# Changes in body shape implicate cuticle stretch in *C. elegans* growth control

**DOI:** 10.1101/2021.04.01.438121

**Authors:** Joy Nyaanga, Christina Goss, Gaotian Zhang, Hannah N. Ahmed, Elliot J. Andersen, Isabella R. Miller, Justine K. Rozenich, Iris L. Swarthout, Jordan A. Vaughn, Niall M. Mangan, Sasha Shirman, Erik C. Andersen

## Abstract

Growth control establishes organism size, requiring mechanisms to sense and adjust growth during development. Studies of single cells revealed that size homeostasis uses distinct control methods. In multicellular organisms, mechanisms that regulate single cell growth must integrate control across organs and tissues during development to generate adult size and shape. We leveraged the roundworm *Caenorhabditis elegans* as a scalable and tractable model to collect precise growth measurements of thousands of individuals, measure feeding behavior, and quantify changes in animal size and shape during a densely sampled developmental time course. As animals transitioned from one developmental stage to the next, we observed changes in body aspect ratio while body volume remained constant. Then, we modeled a physical mechanism by which constraints on cuticle stretch could cause changes in *C. elegans* body shape. The model-predicted shape changes are consistent with those observed in the data. Theoretically, cuticle stretch could be sensed by the animal to initiate larval-stage transitions, providing a means for physical constraints to influence developmental timing and growth rate in *C. elegans*.

**Highlights:**

- Body size measurements of thousands of animals in a dense developmental time course
- Growth rate exhibits nonlinear dynamics in both length and width
- Changes in body shape but not volume occur during periods of increased quiescence
- Dynamics of animal shape consistent with a length-based threshold in cuticle stretch
- Modeling of cuticle stretch dynamics suggests a novel mode for growth control

## 1. Introduction

Growth is a complex process fundamental to development. Individual cells and whole animals must reach an appropriate size to remain competitive in their environment. A larger body size conveys many selective advantages to an organism, including increased success in predation, defense against predation, mating, or intraspecific as well as interspecific competition. Offsetting these advantages, larger organisms require more food resources to grow, take longer to develop, and produce fewer offspring [1]. Therefore, it is critical for multicellular organisms to effectively coordinate the growth of both individual cells and the whole body. Additionally, growth at both of these scales must be coupled with developmental progression to ensure the proper timing of irreversible developmental events.

In recent years, efforts have focused on understanding how organisms control growth to achieve size homeostasis [2–4]. Many of these studies are motivated by the decades-long debate about whether growth is linear or exponential; two separate models each having unique implications for size regulation. In a linear model with constant growth rate, smaller organisms must grow proportionally more than larger organisms to maintain size homeostasis. In this paradigm, organism size can be controlled simply by specifying growth duration. Subsequently, this method of growth control was named the “Timer” model [5,6]. Instead of regulating growth duration, organisms can monitor size and adjust duration of growth to reach an optimal size, often named the “Sizer” model [7–9]. In an exponential model, growth rate is proportional to size. Here, a time-based control mechanism alone would fail to maintain size homeostasis because larger organisms would grow proportionally more during a specified period of time. This difference in growth requires a size-based control mechanism to ensure that growth is halted once a maximum size is reached. Although “Timer” and “Sizer” are the most often proposed size-control models, other models have been suggested. The “Adder” model proposes that a fixed volume is added to a cell or organism during growth [10,11], whereas the “Folder” model specifies that an organism increases in volume by a fixed proportion in order to control growth [12]. It is not trivial to determine which model most accurately describes growth of individual cells or whole organisms because quantitative measurements of growth must be collected at high precision and throughput under tightly controlled experimental conditions. In unicellular organisms, the development of high-throughput experimental techniques in combination with theoretical models have advanced the understanding of size control [13–17]. Progress has been slower for multicellular organisms because cell growth within tissues and tissue growth within organisms often progress at different rates, suggesting that they are likely not regulated in the same ways [18–20].

The nematode *Caenorhabditis elegans* presents both a scalable and tractable multicellular animal model to study growth control. With an adult body length of approximately 1 mm, hundreds of thousands of individuals are easily cultured in controlled laboratory conditions [21]. Moreover, *C. elegans* post-embryonic development is marked by several molts that provide clear developmental milestones [22]. Each molt is initiated by a period of quiescence (lethargus) and terminated once the animal successfully sheds its collagen-rich outer cuticle (ecdysis) [23]. Four molts separate the *C. elegans* life cycle into five distinct stages: four larval stages (L1-L4) and adult. The timing of these molts determines the completion of stage-specific development [24,25] and underscores the importance of growth regulation during *C. elegans* larval development.

A full description of an organism’s development includes the assessment of how growth and body size are regulated. Initial studies of *C. elegans* development described whole-organism growth as a sigmoidal curve characterized by continuous larval growth in length that reaches saturation in adulthood [26]. These early studies hypothesized that molt events had little effect on continuous growth as the *C. elegans* cuticle allowed for stretch during larval stages. Later work determined that larval progression was not continuous but rather piecewise in nature [27]. This study showed that *C. elegans* volumetric growth rate increased from stage to stage such that L1 animals had the slowest rate of growth and L4 animals had the fastest. This finding suggests that *C. elegans* have a mechanism for regulating growth rate, potentially at each molt. Next, researchers using single-animal imaging strategies observed that animals did not advance to the next developmental stage until a critical volume was reached [28]. This finding suggests that *C. elegans* growth follows a “Sizer” model with each molt decision controlled by a volume threshold and further implies that individual cells are able to communicate information about body size to precisely regulate growth. Most recently, live imaging and characterization of body volume heterogeneity revealed that with respect to the start of a larval stage, *C. elegans* volume fold change within a stage is nearly invariant thereby preventing rapid divergence in volume between fast- and slow-growing animals [12]. A mechanism that maintains a constant volume fold change within each larval stage relies on the coupling between growth rate and developmental timing. Notably, such coupling is consistent with recent observations of temporal scaling in *C. elegans* development where despite inter-individual variability in the absolute duration of a larval stage, relative timing of a stage is similar [29,30].

To understand *C. elegans* growth control at the whole-organism level, we used a combination of quantitative measurements and mathematical modeling. We performed a high-resolution longitudinal study of *C. elegans* larval progression and captured high-precision details about animal length, width, volume, and feeding dynamics. By investigating *C. elegans* feeding and growth in tandem for thousands of individual animals, we found decreases in feeding behavior associated with each larval transition that were also correlated in time with changes in growth rate. We used our large-scale measurements of body shape to further analyze the periods of time surrounding each larval transition. At each molt, we observed simultaneous increases in length, decreases in width, and maintenance of volume. Based on the physical characteristics of the cuticle, we propose a “Stretcher” mechanism whereby physical constraints on cuticle stretch influence body shape. We find the Stretcher model-predicted shape changes are consistent with observed data. Animals may be able to detect when the cuticle reaches its maximum capacity for stretch providing a signal to initiate larval-stage transitions.

## 2. Results

### 2.1 Quantitative measurements of *C. elegans* growth

We have optimized a quantitative growth assay that reliably measures small changes in *C. elegans* body size throughout development (Fig. 1). Our method provides both high-throughput and high-precision assessment of developmental growth. In brief, populations of 100,000 animals were cultured in flasks. We cultured six replicate populations of *C. elegans* for a total of 600,000 synchronized and growing animals. Every hour after feeding, a sample of the population from each flask (~300 animals/flask) was collected to measure animal length, width, and feeding rate. The ImageXpress system (Molecular Devices) was used to collect images of sampled animals.Feeding rate, examined using ingestion of fluorescent microspheres as a proxy, and body size were then measured using the COPAS BIOSORT (Union Biometrica). This platform enabled further analysis of life stage and body size, contributing added precision to our measurements.

**Fig. 1.**
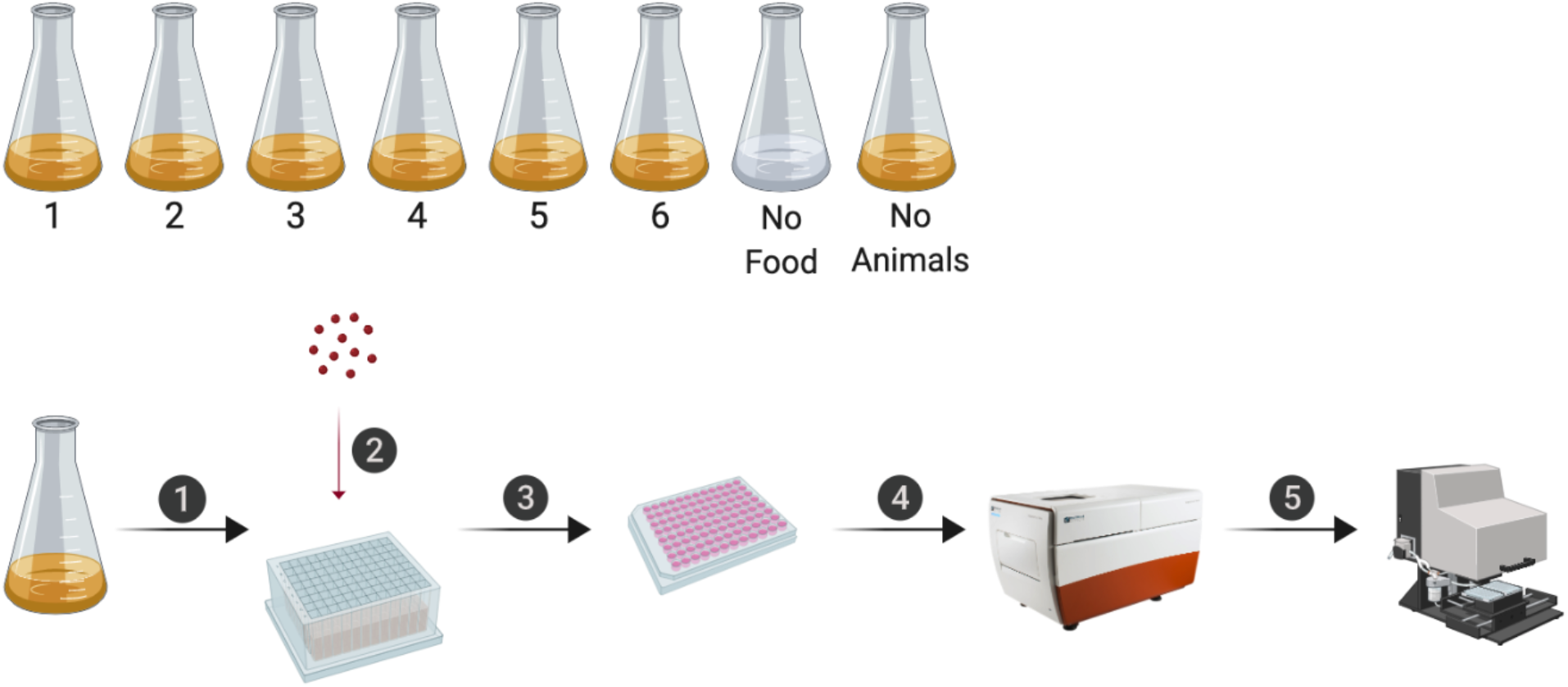
An overview of the quantitative growth assay. Synchronized animals were cultured in flasks where six flasks contained replicate populations of nematodes, one flask had a population of unfed animals, and one flask only contained bacterial food. At each hour of the experiment, all eight flasks were sampled. In step 1, animals were transferred from each flask to a single well of a 96-well microtiter plate. In step 2, fluorescent beads were added to each well. Following a 10-minute incubation period, animals from each well of the deep-well plate were transferred to several wells of a 96-well microtiter plate for step 3. In step 4, animals in each well of the microtiter plate were imaged using the ImageXpress. In step 5, the same animals were measured using the COPAS BIOSORT. This process was repeated every hour after feeding for 72 consecutive hours (see Methods). Schematic of the experimental workflow was created with BioRender.com.

Two measurements of body size were collected from raw data taken from the COPAS BIOSORT: time of flight (TOF) and optical extinction (EXT) (Figure S1). Time of flight is a measurement of body length, and optical extinction corresponds to optical density, a measurement influenced by body length, thickness, and composition [31,32]. We investigated whether optical extinction could be correlated to a different measure of body size using the collection of manual size measurements obtained from images (see Methods). We calculated the median length, width, area, and volume of animals in a subset of imaged wells from each hour of the experiment. We then compared these values to median measurements of animals in each well from the processed COPAS BIOSORT data. We found a strong correlation between manual measurements of animal length from the image analysis and TOF measurements from the COPAS BIOSORT (Figure S2). We also observed an equally strong correlation between manual measurements of animal area and EXT as well as animal width and EXT normalized by body length (norm.EXT). We then calculated animal volume using measurements from the COPAS BIOSORT by using a cylindrical approximation for *C. elegans* shape (see Methods). This result expanded the number of body size parameters that we were able to assess using the COPAS BIOSORT data, allowing us to investigate growth dynamics in length, width, and volume (Fig. 2A-C). To disentangle nematode objects from non-animal objects (bacteria clumps, detritus, shed cuticles), we employed model-based clustering to remove unwanted objects and better examine growth of animals (Figure S3). Lastly, we converted unitless COPAS BIOSORT measurements into microns (see Methods).

**Fig 2.**
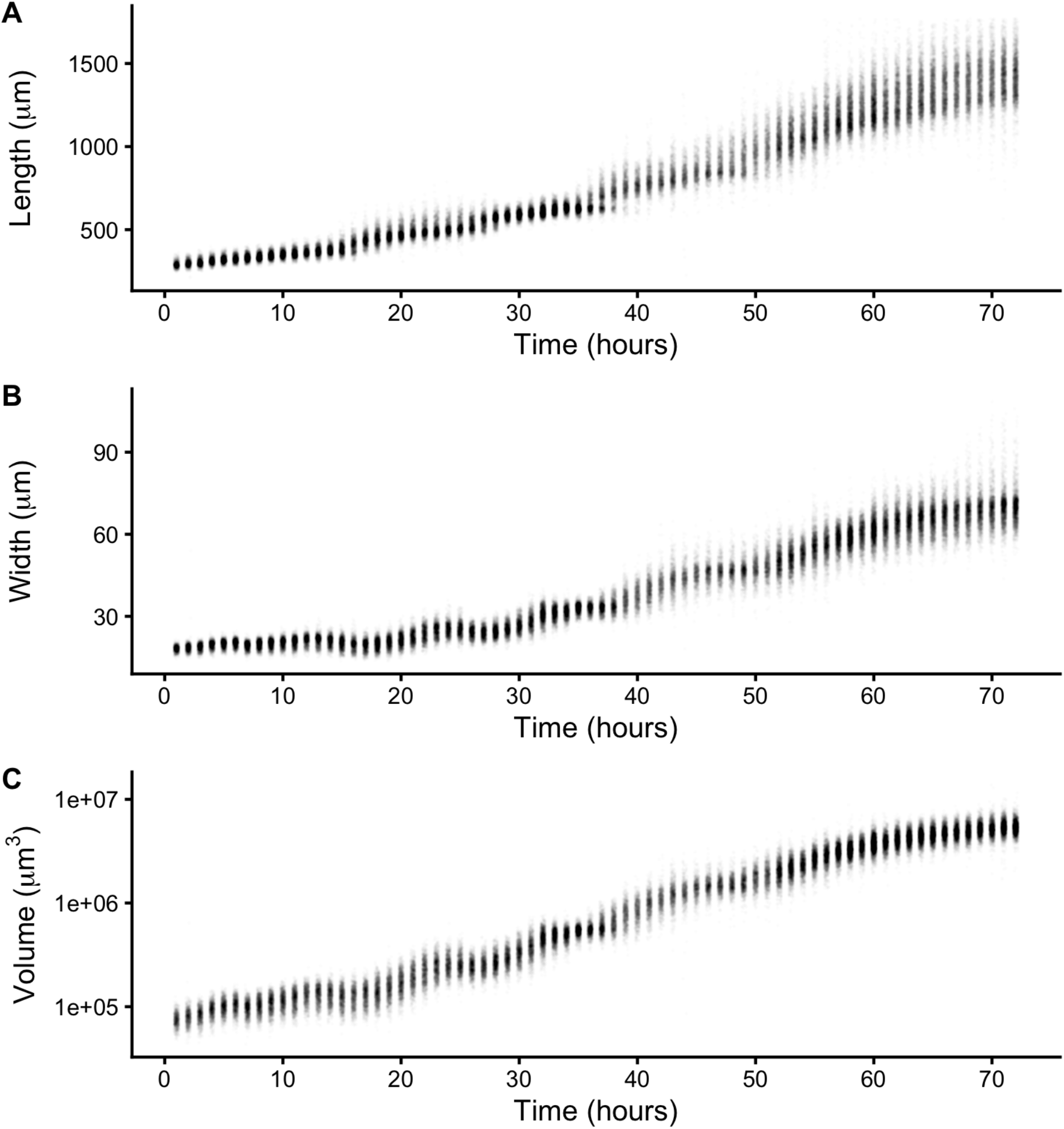
Quantitative measurements of animal size. COPAS BIOSORT data of animal length (A), width (B), and volume (C) after the removal of non-animal objects using model-based clustering methods (see Methods).

We report body length, width, and volume of animals at each hour of development from L1 to adult (Figure S1 and Fig. 2). Historically, growth of *C. elegans* has been shown as a sigmoidal curve where exponential growth during larval stages reaches a maximum rate in adulthood [26]. More recently, researchers have identified that growth curves are discontinuous during development punctuated by larval transitions [27,28]. Using our quantitative growth assay, we captured these small-scale discontinuities in larval growth rate as well as an apparent growth maximum during early adulthood. We noticed that all size variables (length, width, and volume) displayed these dynamics. Objects identified as animals appear to grow in size. However, in particular time windows during development, growth dynamics visibly shift, producing discontinuities in animal growth rate. With these data, we were able to further investigate *C. elegans* growth and size control.

### 2.2 Fluorescence provides a quantitative measurement of animal feeding behavior and developmental progression

In addition to body size and shape, the raw data from the quantitative growth assay included measurements of fluorescence for each animal. To readily assess the thousands of measurements acquired at each hour, we generated summary statistics of median well measurements (Table S1). With these summarized data, we investigated the relationship between feeding behavior and developmental stage. It is well established that temporary suspensions of *C. elegans* feeding occur during each molt[26,33]. As such, active feeding is frequently used to distinguish growing animals from individuals in a molt. We quantified feeding behavior by exposing animals to fluorescent beads the approximate size of bacteria and measuring fluorescence of animals [34]. Because larger animals are able to consume more food and therefore contain more ingested food, we normalized fluorescence by animal area to account for increases in body size (Figure S4). The resulting fluorescence data showed a dynamic pattern (Fig. 3A). At approximately 15 hours, fluorescence steadily increased to a peak before decreasing back to initial levels at approximately hour 27. This pattern, repeated three additional times, established clear time windows of local minimal fluorescence. These local minima represent periods of time where a large proportion of the population had reduced or ceased feeding and therefore suggests time windows where a majority of animals were likely not feeding because they were in a molt. We used a local kernel regression method to estimate a smooth curve and calculate the derivative to identify the time associated with each local minimum (see Methods). We then assessed images collected from the growth assay and found that periods of decreased feeding are concurrent with the presence of shed cuticles, supporting that animals are undergoing a molt during these periods of time (Figure S5). When we overlaid the timing of each local minimum on the population size data, we were able to outline the start and end of each larval stage (Fig. 3B-D). Notably, local minima occurred approximately every ten hours, consistent with well established observations of molt timing [26]. Furthermore, we observed a clear relationship between changes in feeding behavior and growth dynamics where decreases in feeding occurred simultaneously with discontinuous growth in length, width, and volume.

**Fig 3.**
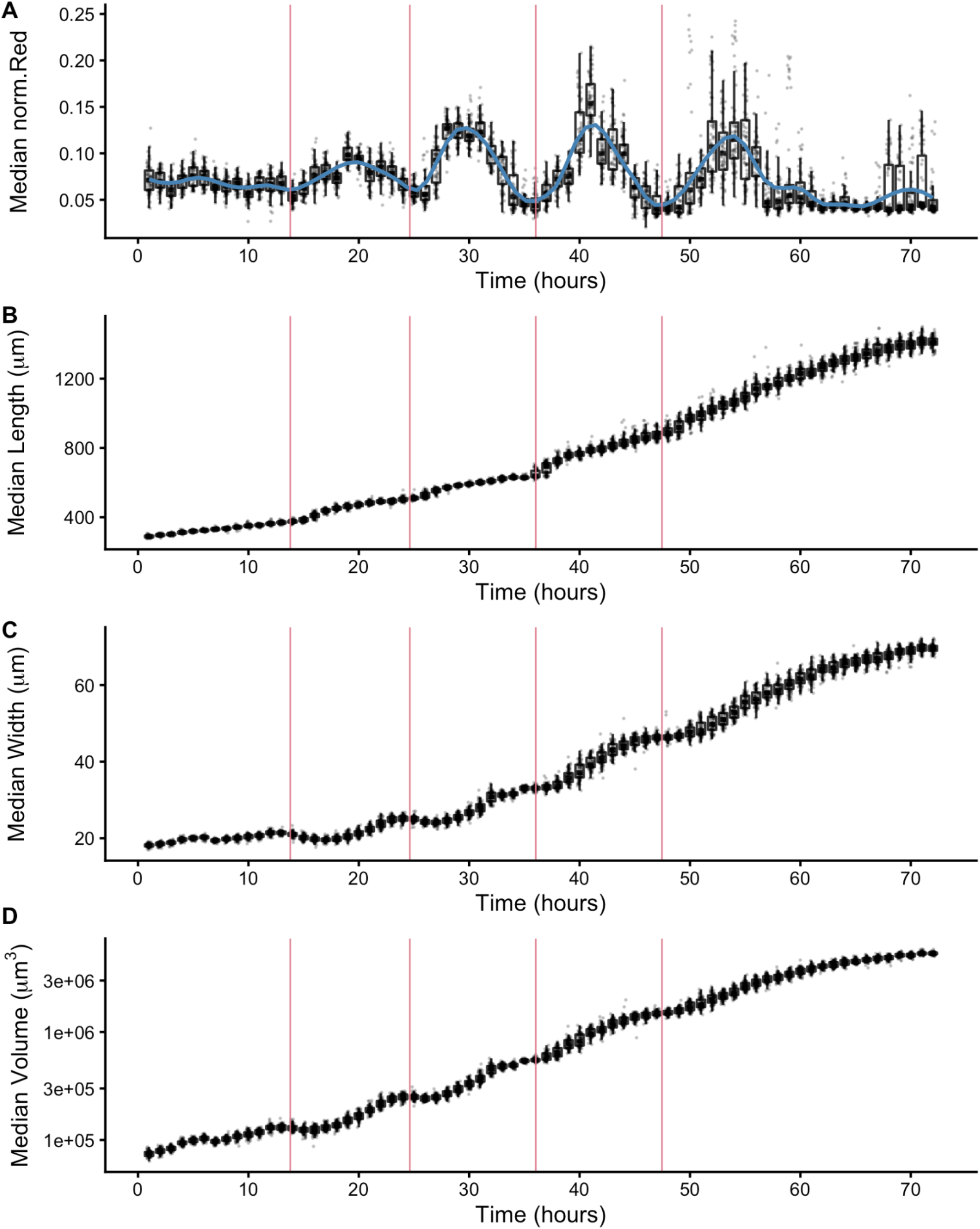
Fluorescence dynamics outline larval stages. (A) Median normalized red fluorescence (y-axis) over time (x-axis) is shown. The blue line represents the kernel regression fit to the data. The red vertical lines correspond to the local minima of the regression and represent the transition between larval stages. Median length (B), median width (C), and median log volume (D) are shown with larval-stage transitions as well. Upper and lower bounds of the box plots correspond to the first and third quartiles. The upper and lower whiskers extend to 1.5 times the value of the interquartile range.

### 2.3 Changes in *C. elegans* body shape occur at larval-stage transitions

Adult body size is ultimately determined by the coordination of developmental progression and rate of growth. To understand how *C. elegans* achieve final size, we must first examine how *C. elegans* grow. Quantitative studies of *C. elegans* growth frequently assess changes in length or volume over time; however, to fully characterize changes associated with growth, it is also important to consider the dynamics of width. Two general models were proposed for *C. elegans* growth in volume: linear and exponential [26–28]. Notably, these volume growth models require different dynamics in length and width. To achieve linear volume growth, length and width must increase at precise sublinear rates that together result in a linear increase in volume. If animal length and width increased at a constant linear rate, then volume would increase at a cubic rate. Alternatively, if both length and width grew exponentially, then volume would fit an exponential model. We sought to identify which model best described *C. elegans* growth behavior but were unable to consistently distinguish between linear, exponential, and cubic models using statistical information criterion because of the similarity in the shapes of the growth curves (Figure S6 and Table S2). This result is not surprising because computational simulations have shown that increases in experimental noise, above 2% added noise, limit the correct identification of growth models [35].

Growth has important implications for how animals regulate size. Size homeostasis requires that growth rate and developmental rate are coordinated. Despite significant variation in individual growth rate, relative timing of *C. elegans* larval transitions is highly similar across individuals [29,30], implying a control mechanism to regulate developmental progression. Early work proposed a size-based growth control model in *C. elegans* [28], although recent work suggests that size homeostasis is achieved through a Folder mechanism [12]. To assess changes in body size and shape during a larval transition, we examined the dynamics of animal length, width, and volume in the hours before, during, and after each molt. We find that for each shape variable, larger animals enter molt first (Fig. 4). We also observe differences in the distributions of lengths during a larval transition compared to widths and volumes. Measurements of animal width and volume remain unimodal throughout a molt, but length does not. As larger animals begin to exit the molt, a rapid increase in body length occurs that leads to the appearance of bimodality of lengths across the population. Importantly, volume remains constant while length increases and width decreases, indicating a change in body aspect ratio not size. Notably, the length increase occurs simultaneously with a decrease in width across the population (Fig. 4D). These changes in the physical dimensions at each larval transition suggests that body shape may be involved in the control of *C. elegans* growth.

**Fig. 4.**
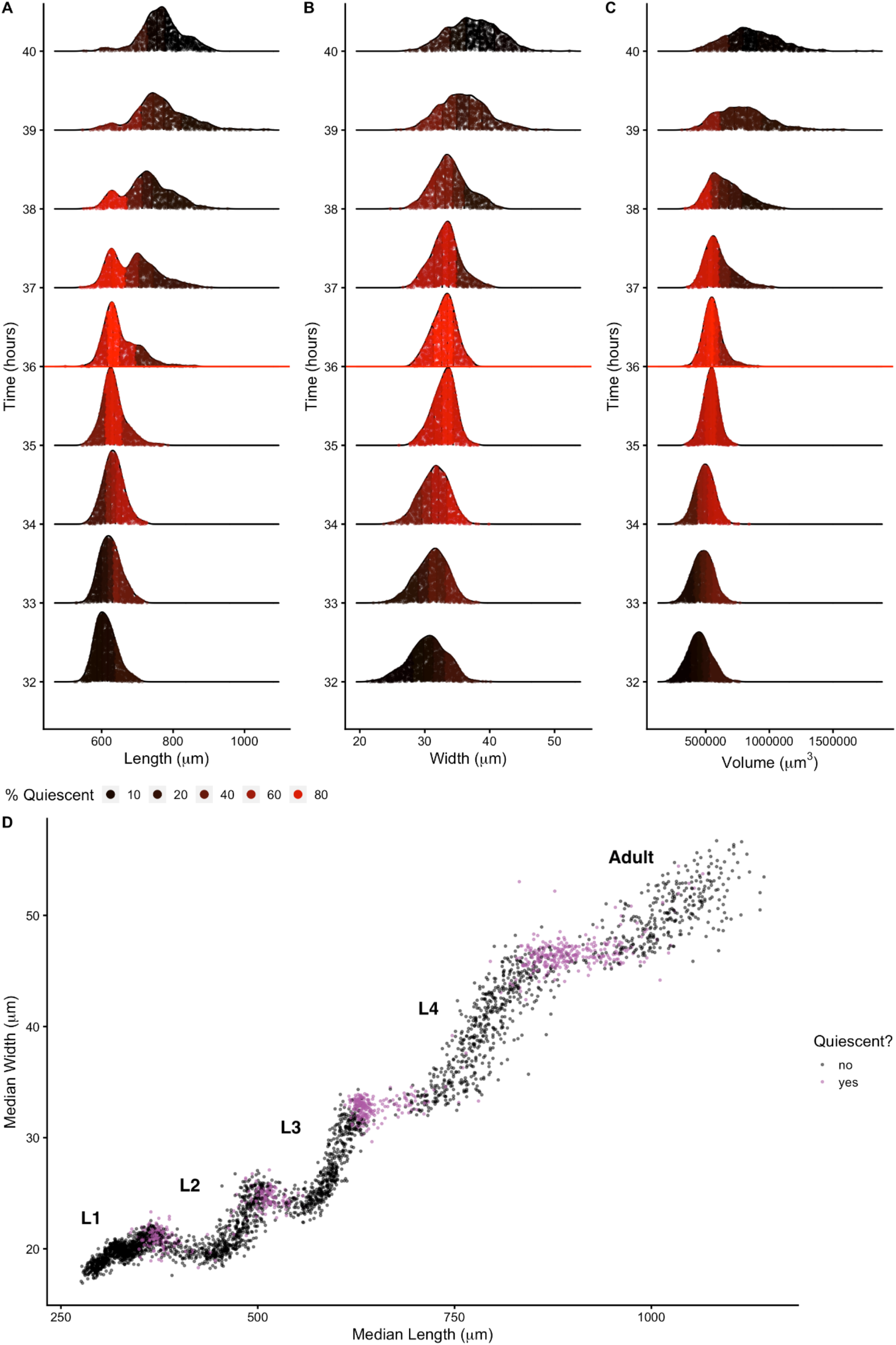
Changes in body shape occur during larval-stage transitions. Population density curves of length (A), width (B) and volume (C) for the hours surrounding the L3 - L4 larval transition (red horizontal line at 36 hours corresponds to the molt). Each distribution was divided into five quantiles. The percentage of quiescent animals present within each quantile was calculated (see Methods), and each quantile was colored to reflect this percentage. In all shape variables, quantiles that contain the largest animals displayed an increase in quiescence earlier than quantiles that contain the smallest animals. These dynamics were consistent across all larval-stage transitions (Figure S7). (D) Median width vs. median length for experimental hours 1 - 55. Purple indicates measurements that fall above the quiescence threshold (see Methods). Simultaneous changes in length and width occur during periods of increased quiescence.

### 2.4 Modeling *C. elegans* cuticle stretch dynamics

#### 2.4.1 Cuticle stretch could be a sensor for larval-stage transitions

Previous studies theorized that the internal mechanism for sensing body size and triggering molts in *C. elegans* is driven, in part, by the properties of the collagen-rich cuticle [12,28]. Many cuticle collagen mutations cause morphological defects in nematode shape [36]. Some of these mutants are shorter than the wild-type but do not have differences in animal width, implying that the cuticle affects length and width independently [37]. The *C. elegans* cuticle does not grow through the addition of new material but rather stretches to accommodate increases in animal body size. Cuticle stretch is likely limited by the material properties of the cuticle. The *C. elegans* cuticle is primarily made of cross-linked collagens organized into lateral ridges and circumferential bands [38]. Commonly found in many biological systems, collagen-based materials are fairly flexible under low stress conditions. However, as stress increases, collagen fibrils become elongated and orient in the load bearing direction leading to a decrease in elasticity [39]. Previous work using atomic force microscopy revealed a high level of biomechanical stiffness at the circumferential bands [40], leading others to speculate that mechanical strain on these structures is likely adjusted as internal body pressure changes [41]. Additionally, in *nekl-3(sv3)* molting mutants, the cuticle is not properly removed from the middle part of their body, leaving the free head and tail to grow normally while the encased middle is constricted by the old cuticle to pre-molt dimensions [42]. Given this body restriction, we speculated that the old cuticle stretches beyond its tolerance, becomes stiff, and constricts the center of the nematode relative to the growing head and tail size. To our knowledge, it is unknown if the cuticle stretches enough during the course of normal development to become stiff in either the length or width directions. If the cuticle does become stiff before a molt, *C. elegans* may be able to sense the reduction of elasticity or “stretchiness,” and use this signal, along with others, to determine when to initiate a molt. To detect possible changes in cuticle stretch during normal development, we asked how changes in cuticle elasticity will impact body shape.

To understand the impact of cuticle elasticity on body shape, we developed a “Stretcher” model independent of our measurements during development. We propose that the nematode passes through three distinct regimes related to cuticle stretch: linear stretch dynamics, non-linear stretch dynamics, and the larval-stage transition (Fig. 5). These regimes arise naturally from the following biologically supported assumptions. The cuticle structure is anisotropic, possibly leading to distinct stiffness properties in the length and width directions [43,44]. We approximated the cuticle as a hollow cylinder of negligible thickness filled by the body of the nematode. Growth was modelled as internal pressure evenly applied to the cuticle in all directions. An anisotropic, elastic cuticle responds differently to pressure from growth during linear stretch, nonlinear stretch, and post-molt relaxation, leading to differences in shape during each stage of development.

**Fig. 5.**
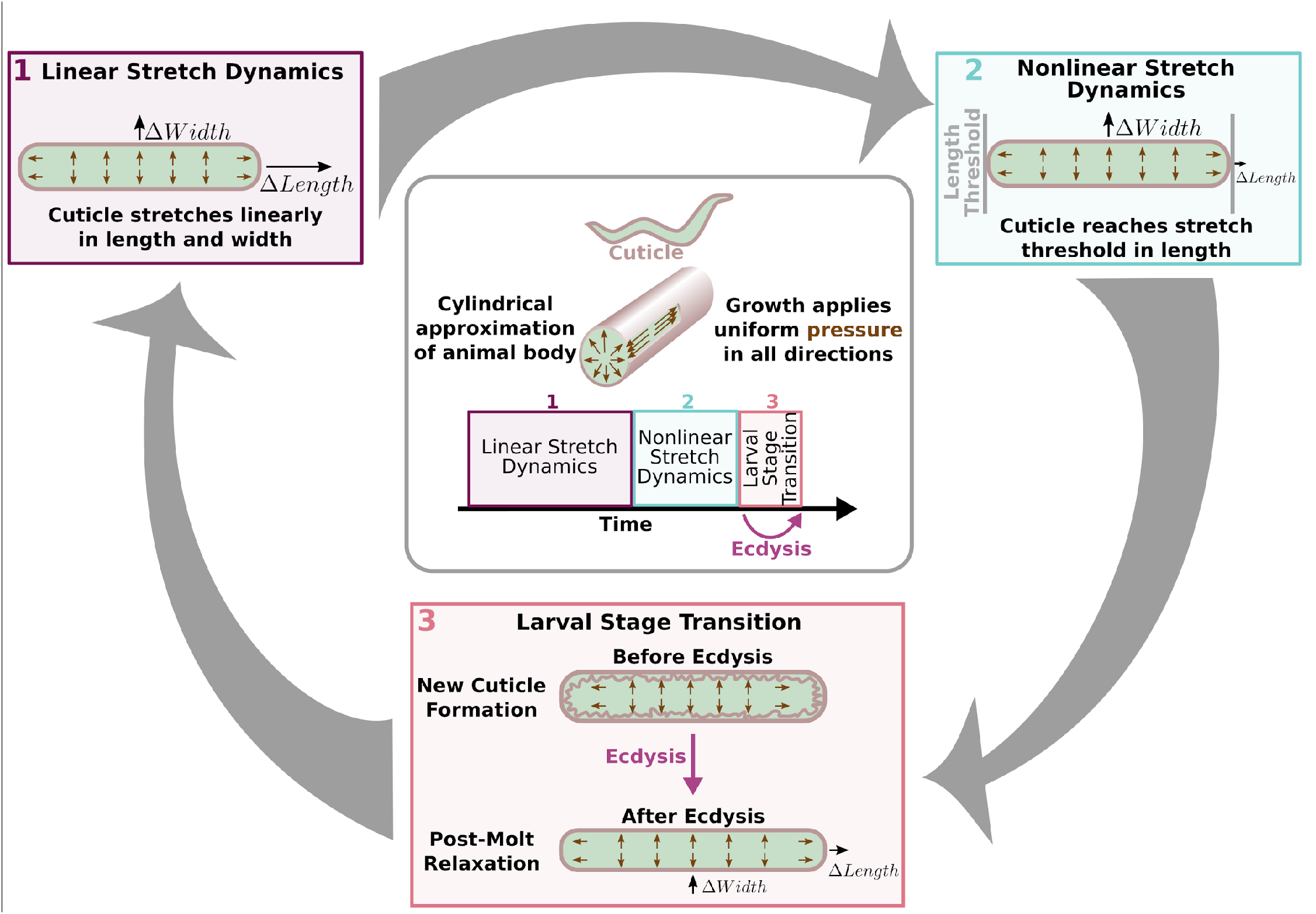
Cuticle stretch dynamics guide larval-stage transitions. The Stretcher model describes each larval stage as a cycle. Nematodes are modeled as a cylindrical object with a thin cuticle epidermis. (Box 1) Linear Stretch Dynamics: uniform growth pressure stretches the cuticle linearly in both length and width. (Box 2) Nonlinear Stretch Dynamics: the cuticle has reached a stretch threshold in length, and under uniform growth pressure the length stretches less (sub-linear) and width stretches linearly. (Box 3) Larval Stage Transition: a new cuticle is formed and the old cuticle is shed (ecdysis), removing constraints in length. The nematode body “relaxes” in length, causing an increase in length, a decrease in width, and constant volume.

In the linear stretch regime (Fig. 5), the cuticle would be linearly elastic in both the length and width directions, stretching proportionally to the pressure exerted on the cuticle. Previous work found evidence for a linearly elastic cuticle [45,46] in animals expanded in a negative external pressure environment or after positive force was applied to the cuticle. Gilpin *et al*. found evidence of linear elasticity in the nematode body. We conjecture that this linear elasticity is caused by the constraints applied by the cuticle [45,46]. A linearly elastic cuticle will have Δ*L* stretch in the length direction and Δ*W* stretch in the width direction, each related to growth-applied pressure Δ*p* by

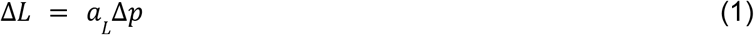

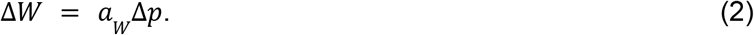

The “stretch coefficients” in length, *a_L_* and width, *a_W_*, measure the stiffness of the cuticle (File S5, Eq. S4-S13). Smaller values correspond to a stiffer material, which is less able to stretch in response to pressure. The stretch coefficients are constant in the linearly elastic regime and are determined by geometric constants and material properties. The ratio of the change in length (Eq. 1) and width (Eq. 2) produces a pressure-independent relationship that depends only on the ratio of the geometric and material properties, which can be verified using measurements of length and width (Fig. 3). During the linearly elastic regime, the ratio of growth in width to growth in length is constant throughout a larval stage where the cuticle properties are fixed as in

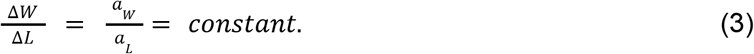

In the non-linear stretch regime (Fig. 5), growth continues to apply pressure to the cuticle uniformly in all directions. As observed in *nekl-3(sv3)* mutants, the cuticle can restrict body growth [42]. Once outside of the linearly elastic regime, the cuticle would hardly stretch, even under large forces. We hypothesize that this shift from linear to nonlinear regimes can provide a mechanism for size-sensing and cues the larval-stage transition (Fig. 5). In principle, this transition could occur in either the width or length directions. For simplicity, we illustrate a transition from linear to non-linear stretch in the length direction while linear stretch in the width direction is maintained. In the nonlinear regime, the stretch in the length direction in response to pressure becomes

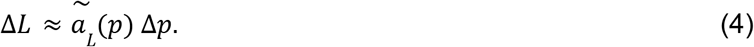

The nonlinear “stretch coefficient,” 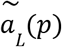, is no longer constant and decreases with increasing pressure. It is smaller than *a_L_* because the cuticle has become less elastic than in the linear regime. If the length-direction enters the nonlinear regime and has reduced stretch response and width has the same constant stretch response, then we expect the Δ*W*/Δ*L* ratio to increase

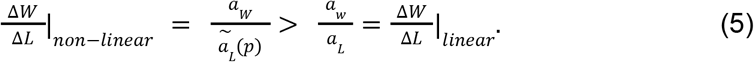

If the width-direction enters the nonlinear regime first, then the Δ*W*/Δ*L* would decrease. It is possible that both length and width could enter the nonlinear regime at around the same time, in which case the change in the Δ*W*/Δ*L* ratio would depend on the differences in the pressure-dependent stretch coefficients. This prediction motivates an analysis of the Δ*W*/Δ*L* ratio of the shape data (Fig. 4), to determine if any sign of a nonlinear stretch regime in either direction is observed.

During the larval-stage transition (Fig. 5), a new, larger cuticle is formed beneath the old cuticle that is shed during ecdysis. Because the old cuticle constrained growth in length, we predict a rapid increase in the length direction when the constraint is removed. Animal volume is conserved as growth does not occur during this process. Therefore, the relaxation in length is accompanied by a corresponding decrease in width.

#### 2.4.2 Discontinuities in animal growth rate might be driven by limits on cuticle stretch

To test whether the shape dynamics predicted by the Stretcher model are consistent with the data, we analyzed the relationship between measured nematode length and width over developmental time. This analysis requires no model fitting or assumptions about which direction may enter a nonlinear regime first, and it involves only calculating Δ*W*/Δ*L*. Although the shape relaxation for an individual animal is expected to happen at a much shorter time scale than data collection (seconds to minutes), the measured shape change within the population occurs on a larger time scale (several hours). As such, we can observe the sudden shape change at transitions by assessing changes in length and width of the population (Fig. 6). Doing so, we detected all three regimes predicted by the Stretcher model in the COPAS BIOSORT data: linear stretch, non-linear stretch, and relaxation (Fig. 6A). In all larval stages, we observed an approximately constant instantaneous Δ*W*/Δ*L* ratio, consistent with a linear stretch regime (Fig. 6B and Fig. 6C). We also detected a large slope decrease during the L1 stage, which could correspond in time to the metabolic decision for entry into the dauer stage [47] or divisions of the seam cells [48]. Although transitions in the slope (Δ*W*/Δ*L* ratio) are difficult to detect at all larval stages because of noise amplification and population effects (Figure S9), we observed a sharp slope increase, consistent with a non-linear stretch regime in length prior to lethargus, near the end of L2 and L3 stages. Following the nonlinear stretch regime, we noted a simultaneous increase in length and decrease in width at the transition between larval stages, consistent with a length threshold in the Stretcher model (Fig. 5D, Fig. 6A, and Figure S8). As a whole, the changes in animal shape are consistent with the hypothesis that the cuticle reaches the nonlinear elastic regime before larval-stage transitions during normal development. The ability to sense when a critical cuticle stiffness is reached would allow animals to monitor fold changes in body growth within each larval stage, serving as a connection between growth rate and developmental timing.

**Fig. 6.**
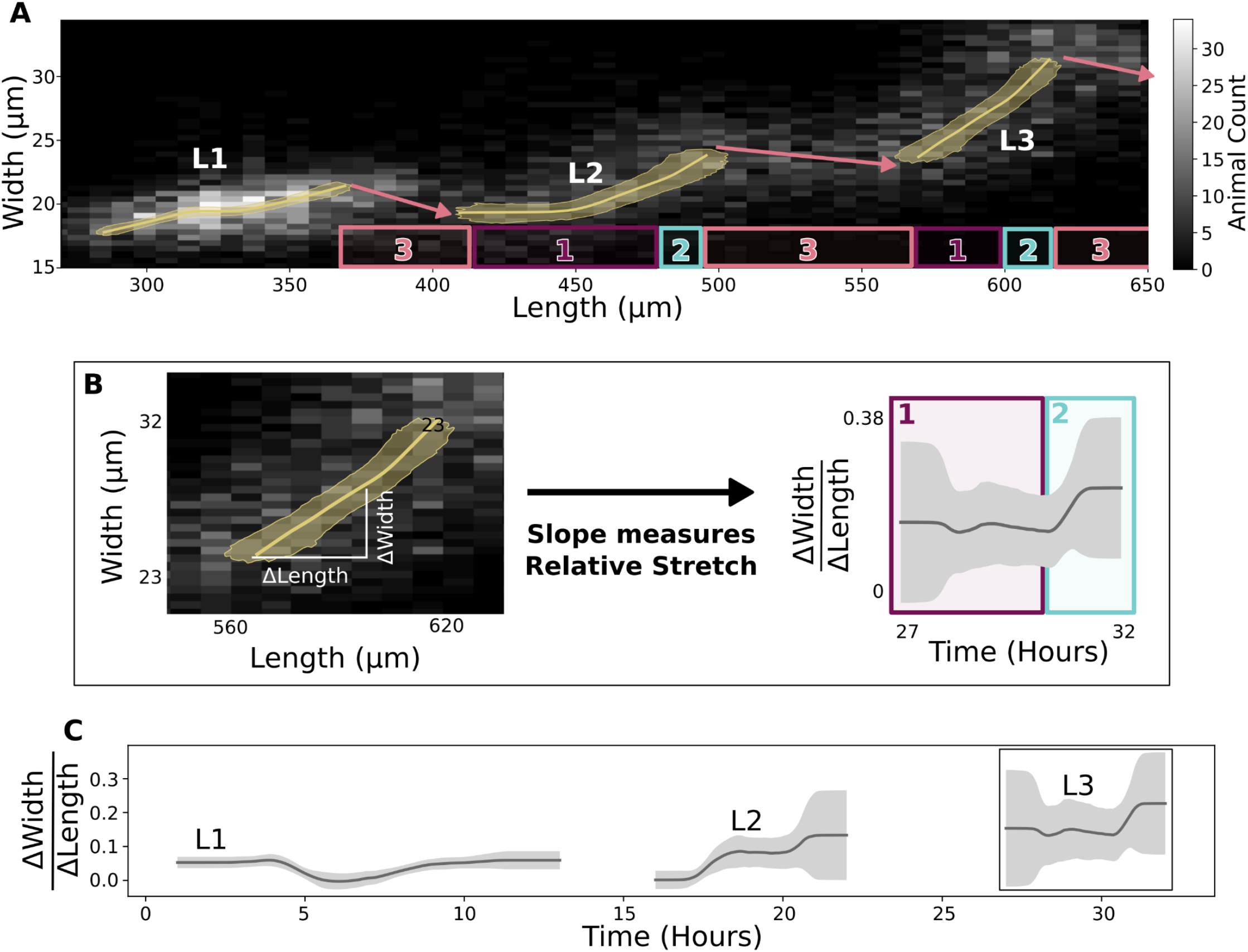
Experimental data are consistent with a length threshold in cuticle stretch. (A) A grayscale histogram of the width (y-axis) vs length (x-axis) of all sampled animals in replicate 2. The range of all bootstrap regressions is in gold. (B) Demonstration of calculating the ratio of width-to-length stretch as the local slope using L3. Left panel is a repetition of L3 data from Fig. 6A. Right panel is a repetition of results from Fig. 6C. (C) Within a larval stage, the ratio of width to length stretch varies over time. The standard deviation captures population variation (grey) (see Methods).

## 3. Discussion and conclusions

Using an integrated image-based and flow-based phenotyping strategy to precisely evaluate feeding, growth, and molt dynamics at high replication, we detected oscillations in feeding behavior consistent with larval progressions and used these dynamics to define larval stages. We observed changes in body shape at each larval-stage transition that can be driven by differences in physical cuticle properties along length or width (anisotropy). These results suggest a mechanism by which animals sense their size to trigger larval-stage transitions by detecting the physical stretch of the cuticle, demonstrating how physical constraints can influence developmental timing and growth rate.

### 3.1 A potential role for cuticle stretch in the timing of larval-stage transitions

Measurement of both animal length and width allowed us to observe changes in body shape as well as body size. We propose a mechanism in which a stretch threshold along the body length axis acts as a trigger to larval-stage transitions. Previously, a Folder mechanism for *C. elegans* growth has been suggested [12]. Mechanical stretch sensing could provide organisms a way to couple the rate of growth and development to maintain a constant volume fold change within a larval stage. In this way, smaller animals would reach a stretch limit at a smaller size as the cuticle would only stretch a percentage of its original size before reaching a threshold. For cuticle stretch to trigger larval-stage transitions, animals must either have the ability to measure the amount the cuticle has stretched or the stiffness of the cuticle. Across biological systems, cells can respond to the stiffness of their environment using mechanosensitive components [49,50], but few examples in tissues or whole-organisms are known. In *C. elegans*, it has been demonstrated that hemidesmosomes, which connect the cuticle and the epidermis, are mechanosensitive during embryogenesis [51,52]. Additionally, dense bodies, which connect the epidermis and muscles, are hypothesized to be mechanosensitive as well [25,53–55]. Changes in cuticle composition, and presumably stiffness, have been shown to also affect well known growth controlling pathways such as the BMP signaling pathway [56]. These possible mechanosensitive components could monitor the stiffness of the cuticle and be part of the signaling pathways that regulate the initiation of molts. Further experiments are required to explicitly test whether these components control larval-stage transitions but disentangling epistatic growth effects in mutants from specific mechanosensation effects might be difficult. It is also important to note that evidence for a stretch-based mechanism for growth control does not preclude the possibility of a developmental timer. It is likely that physical constraint-based events are part of a larger regulatory system coordinating developmental decisions.

Our analysis of Δ*W*/Δ*L* ratio changes over larval stages provides a first approximation of the timing of cuticle stretch properties. The increase we observed in the Δ*W*/Δ*L* ratio, followed by sudden relaxation to a longer skinnier animal is consistent with a decrease in the elasticity of the cuticle in the length direction before a molt. These observations of shape change also indicate that *C. elegans* do not preempt the shape change by molting before they hit the “stretch limit”, meaning that the decrease in elasticity could be detected by the animal and this stretch-based physical constraint mechanism could play a role in developmental decisions. Interestingly, when observing the L4 to adult transition, others have detected anisotropic constriction on the transverse (width) axis followed by gradual relaxation driven by rearrangements in cortical actin networks [57]. Single-worm, high frequency measurements targeting hours surrounding the sudden width-to-length ratio increase, are needed to better resolve cuticle shape dynamics. Although analysis of the larval growth dynamics for *C. elegans* body shape mutants (*dpy*, *lon*, or *sma*) may provide insights into Δ*W*/Δ*L* ratio variation, measurements of whole-animal length and width only provide a total stiffness estimate, leaving us unable to distinguish the contributions of cuticle stiffness from other tissues. For example, previous work has shown that bodies of *dpy-10(e128)* mutants are twice as stiff as the wild type [58], and *dpy-5(e61)* mutants are softer than wild-type animals [45]. Authors have speculated that this difference in stiffness is caused by an increased internal glycerol concentration in *dpy-10(e128)* animals that is absent in *dpy-5(e61)* animals [58]. Therefore, although these mutations impact body shape similarly, they do not have the same effect on body stiffness, indicating that body shape alone does not predict body stiffness. To assess cuticle properties independent of other nematode tissues and organs, future experiments probing the stiffness of free cuticles are necessary.

Additionally, within the L1 stage, the relative stretch measured in width and length did not follow the pattern observed in other larval stages. We observed a mid-stage dip in the width-to-length ratio that is otherwise approximately constant throughout the L1 stage. Experiments exploring the structural properties of cuticles at all larval stages might help to determine where the L1 shape changes originate.

### 3.2 Development comprises complex interactions of growth regulation across diverse scales

Our results demonstrate that *C. elegans* may use physical constraints on animal size, in tandem with other regulatory mechanisms, to control growth rate and determine developmental transitions. This type of regulation could be applicable to organisms with stiff cuticles or other physical barriers to growth, like many species of Ecdysozoa. The control of whole-organism growth requires cells, tissues, and organs to orchestrate final size and cell number. In *C. elegans*, cell number is precisely defined and invariant from animal to animal [59], so the final adult size of an individual must come from cell size as opposed to number. Future studies should focus on how whole-organism size is determined by the integration of cell, tissue, and organ size. By incorporating these different developmental scales, the Stretcher model can be refined to completely describe how physical constraints on parts of the organism impact the whole. *C. elegans* gives investigators a powerful system to investigate animal-to-animal variation in developmental trajectories across each of these scales.

## 4. Materials and methods

### 4.1 Worm culture

The canonical laboratory strain N2 was obtained from the *C. elegans* Natural Diversity Resource [60]. Animals were cultured at 20°C on 6 cm plates of modified nematode growth media (NGMA)[61], which contained 1% agar and 0.7% agarose seeded with *E. coli* OP50 bacteria.

### 4.2 Bacterial food

*E. coli* HB101 bacteria were prepared from cultures grown for 15 hours in Superbroth and then pelleted by centrifugation. HB101 bacteria were diluted to OD100 in K medium (51 mM NaCl, 32 mM KCl, 3 mM CaCl2, and 3 mM MgSO4 in distilled water) and stored at −80°C. Bacteria were thawed and fed to animals at a concentration sufficient to sustain population growth from hatching to adulthood (OD20).

### 4.3 Growth of the animals

Populations of animals were propagated on NGMA plates for two generations without starvation. In the third generation, gravid adults were bleach-synchronized [62]. Embryos were resuspended in K medium, aliquoted into a 500 mL flask at a concentration of one embryo per μL, and allowed to hatch overnight. The following day, arrested L1s were fed HB101 bacteria at a final concentration of OD20 in a final flask volume of 100 mL K medium and HB101 food. Animals were grown for three days at 20°C with constant shaking. Following these three days, adult animals were bleach-synchronized once more and embryos were aliquoted to seven replicate 500 mL flasks at a concentration of one embryo per μL in 100 mL. The following morning, six flasks were fed HB101 bacterial food at a final concentration of OD20 in a final flask volume of 100 mL K medium and HB101 food. Two additional flasks were included to control for L1 animal size and possible clumping of bacterial food: one flask contained L1 larvae but did not have food added and one flask contained no larvae but the same concentration of HB101 bacteria as the six flasks containing L1 larvae. All replicate flasks were kept in an incubator at 20°C with shaking for the duration of the experiment. A small temperature gradient of 1.25°C was recorded in the shaking incubator with the highest temperature readings on the right side and lowest temperature readings on the left side (S1 File). This slight variation in temperature contributed to variation in developmental rate among replicates based on position within the incubator (replicates were placed in numerical order with replicate 1 positioned on the far right side of the incubator).

### 4.4 High-throughput measurements of body size and fluorescence

Flasks were sampled each hour beginning one hour after feeding and continuing for 72 consecutive hours. At each hour, 500 μL was removed from each flask and transferred to a well of a deep 96-well plate. Each flask was sampled at each time point. Fluorescent polychromatic beads (Polysciences, 19507-5) with a 0.5 μm particle size were added to each well at a final concentration of 3.64×10^8^ beads/mL and incubated at 20°C for 10 minutes with shaking. Following the bead incubation, 30 μL from each well of the deep 96-well plate was aliquoted to a 96-well microtiter plate. The process was repeated 11 times to 11 separate wells of the same microtiter plate with pipetting to mix the well contents from the deep 96-well plate. Animals were then treated with sodium azide at a final concentration of 50 mM to paralyze and prevent defecation of the ingested beads. The 96-well plate was imaged with an ImageXpress Nano (Molecular Devices, SanJose, CA) using both 2x (Nikon MRD00025) and 10x (Nikon MRH00101) objectives. The ImageXpress Nano acquires brightfield images using a 4.7 megaPixel CMOS camera. Images are stored in 16-bit TIFF format. Finally, animals were scored using a large-particle flow cytometer (COPAS BIOSORT, Union Biometrica, Holliston MA). The COPAS BIOSORT sheath flow rate was kept at a constant 10.3 ±0.1 mL per minute to reduce variability in length measurements.

### 4.5 Image processing

Manual measurements of animal size were performed using the free Java image-processing program ImageJ [63]. Well images for the six replicate flasks, excluding controls, were loaded into ImageJ software. Length was measured from head to tail, and width was measured at the widest point of the animal. Five animals were measured per well across thirty total wells for each hour. Measurements were repeated for all 72 time points in the assay. Body length and width were used to estimate cross-sectional area (length*width). This metric was used to describe animal area for the extent of the text. Volume was calculated from body length and width by approximating the animal as a cylinder. Pixels were converted to μm using a conversion factor of 3.2937 pixels/μm.

### 4.6 Data processing

The COPAS BIOSORT was used to collect measurements of animal length (TOF), optical extinction (EXT), and fluorescence for every animal in each well. These traits measure properties of nematode development and, as such, increase as animals progress to adulthood [31]. Optical extinction measurements correspond to the amount of light absorbed over the full length of an animal as it passes through the instrument. An additional measurement (norm.EXT) can be calculated by normalizing optical extinction by length. The raw data collected were imported and processed using the *easysorter* R package [64].

The COPAS BIOSORT data were analyzed further using Gaussian finite mixture modeling as implemented in the *mclust* R package [65]. These probabilistic models assume that data are generated from a mixture of multivariate normal distributions and, therefore, can be used to classify unstructured data into meaningful groups. Specifically, the *mclust* package fits a mixture model to data and selects the optimal number of clusters using an expectation-maximization algorithm and Bayes Information Criteria. For model-based clustering, log transformed animal length (logTOF) and log transformed optical extinction (logEXT) were used as inputs for the *Mclust* function. Data from each hour of the experiment was analyzed by replicate and clusters that did not appear to include majority animal objects were identified and removed as described previously [66]. This processing removed non-animal objects such as bacterial clumps, shed cuticles, and next generation larval animals from the time-course data.

We used a numpy polyfit regression of well-median data from the COPAS BIOSORT and image measurements to convert TOF and norm.EXT data to microns (File S5, Eq. S1-S3). Only the unit-corrected BIOSORT data were used for further analysis.

### 4.7 Molt analysis

Fluorescence data obtained from the COPAS BIOSORT was used as a proxy for feeding behavior to distinguish animals in a molt from growing animals. First, fluorescence was normalized by EXT to account for the ability of larger animals to consume more food and beads. Next, an analysis of variance statistical model was fit to the fluorescence data normalized by EXT to determine the amount of variance contributed by replicate and well (Table S1). A local kernel regression smoothing method was then applied to the residuals of the variance analysis using the *lokern* R package [67]. Residuals were used to address only the differences over time and ignore minor variation among replicates and wells. The local minima of the regression function were found by solving for where the first derivative of this function equaled zero. The time associated with each local minimum was used to represent the timing of each molt. Molts occurred at 14, 25, 36, and 48 hours.

To identify periods of time that contained a majority of growing animals, the inflection points of the regression function were calculated by solving for where the second derivative of the function equaled zero. Time points between inflection points that did not contain a local fluorescence minimum were considered as growth periods. These hours were 1-13, 17-22, 27-32, and 39-45 corresponding to L1, L2, L3, and L4 growth periods.

Each molt is initiated when animals enter lethargus: a behavioral state where animals cease active feeding. To classify individual animals as in a molt or growing, we set a quiescence threshold using fluorescence measurements at each local minimum. The fluorescence measurement at each local minimum was as follows: 0.07, 0.06, 0.06, 0.06. The average of these measurements (0.06) was used as the fluorescence threshold signifying quiescent behavior. Any individual animals that fell below this threshold fluorescence value were designated as in a molt and animals above this threshold value were classified as growing.

### 4.8 Comparison of model fits

To determine the volume growth model, we fit linear, exponential, and cubic functions to data designated as growth periods for each larval stage. Both linear and nonlinear functions were fitted using least-squares regression. Akaike’s information criterion (AIC) [68] and Bayesian information criterion (BIC) [69] were goodness of fit criteria used to evaluate candidate models. To assess the strength of evidence for each candidate model, we identified the model with the smallest AIC/BIC value and assessed the difference between this value and the AIC/BIC of the other two models. The magnitude of the difference was used to determine the level of support for each candidate model as previously described [70,71]. All model fits and analysis were performed using the *stats* R package.

### 4.9 Stretcher model analysis

To analyze shape dynamics, length and width data from growth time periods were extracted from the full COPAS BIOSORT population data and analyzed from each replicate separately to avoid issues with replicate variability. For replicate 2, the hours defining growth periods were 1-13, 16.37-22.39, and 26.93-32.96; corresponding to L1, L2, and L3. Hours defining larval stages were rounded as data was collected at exact hour increments. The L4 stage was excluded from the analysis because of the high variability within the population. We applied a local kernel regression, *lokern* R package [72], to smooth the population dynamics of length and width. To calculate mean and standard deviation, the smoothed population measurements were bootstrapped using 2,000 samples with replacement (File S5, Algorithm S1). To determine cuticle properties throughout larval stages, we calculated the mean ratio of derivatives of regression width and length. Error for this ratio was calculated using error propagation to pass the bootstrap variation through the ratio (File S5, Eq. S17-18).

## Data availability

The authors state that all data necessary to confirm the conclusions of this work are within the text, figures, and supplementary material. All files and code for analysis and generation of figures and tables are archived on GitHub (https://github.com/AndersenLab/C.elegans-growth-manuscript).

## Author contributions

E.C.A and N.M.M conceived the project. J.N., G.Z., and E.C.A designed, optimized, and performed the experiments. J.N., H.N.A, E.J.A, I.R.M, J.K.R, I.L.S., and J.A.V. collected manual size measurements of animals from images. J.N. processed the COPAS BIOSORT data to remove non-animal objects. J.N, C.G., and S.S. analyzed the results. C.G. and S.S. developed the theory and tested the models. E.C.A and N.M.M supervised the research and the development of the manuscript. J.N, C.G, and S.S. wrote the first draft of the manuscript; J.N., C.G., G.Z, N.M.M, S.S., and E.C.A. edited the manuscript.

## Acknowledgements

We thank Nicole Roberto for help with high-throughput sampling. We thank Jiping Wang and Keren Li for helpful advice about statistical data analysis. We would also like to thank members of the Andersen group and the Mangan group for their helpful comments on the manuscript.

