## Supplement for "Changes in body shape implicate cuticle stretch in *C. elegans* growth control"

#### Supplementary material

**S1 File. Incubator temperature data.** Temperature recordings of each position within the shaking incubator used for the growth experiment. (CSV)
**S2 File. COPAS BIOSORT growth data.** Raw growth data collected from the COPAS BIOSORT and processed using the *easysorter* R package to compile information from each well. (CSV)
**S3 File. Pruned COPAS BIOSORT growth data.** Processed data from the COPAS BIOSORT following implementation of the *mclust* R package and removal of clusters containing non-animal objects. (CSV)
**S4 File. Image growth data.** Manual measurements of animal size acquired from images. (CSV)
**S5 File. Model derivations.**

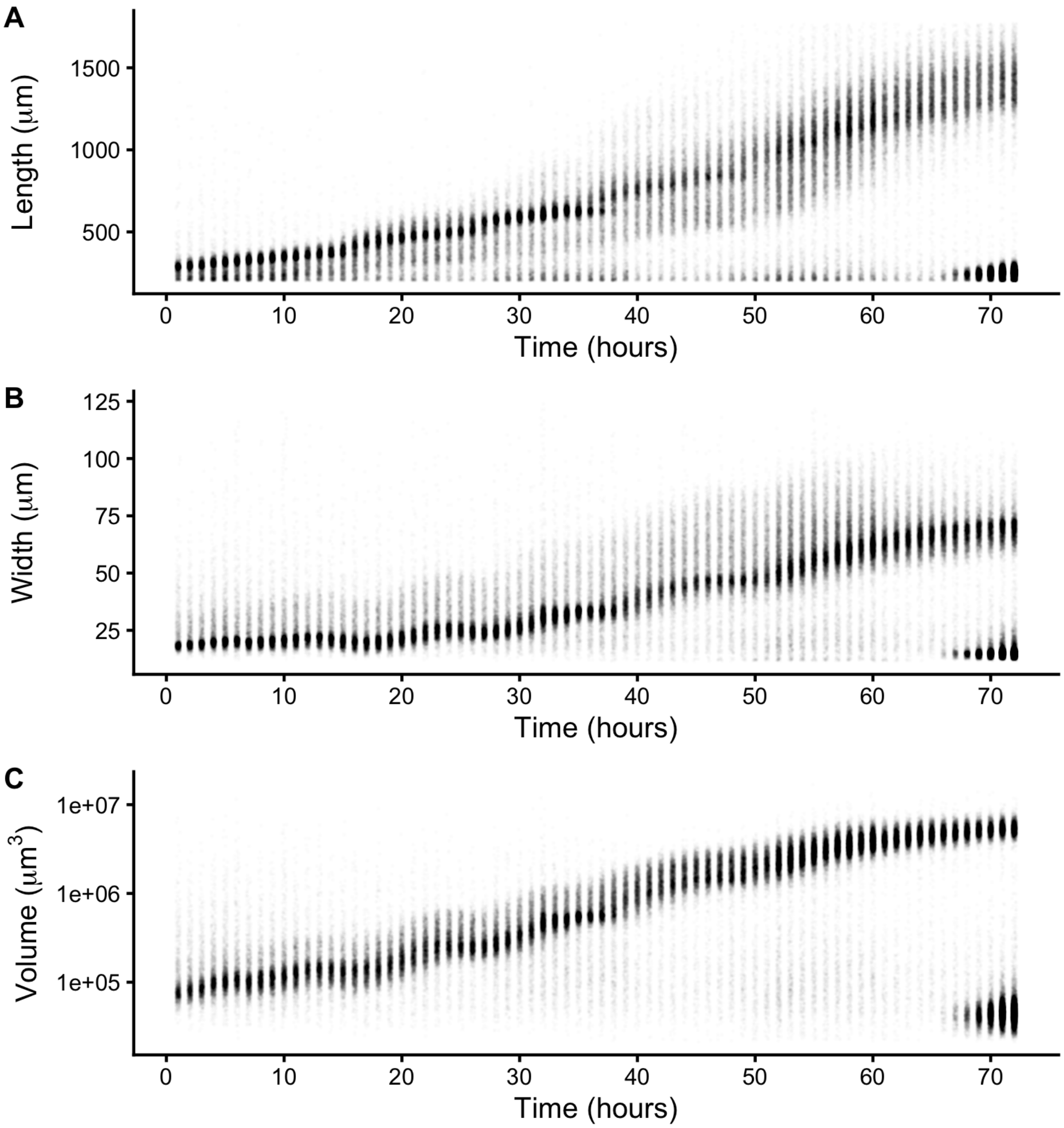


**Figure S1. Raw measurements of animal size.** Raw COPAS BIOSORT data of animal length (A), width (B), and volume (C) are shown here. After 60 hours, animals have developed to the adult stage. Smaller objects observed after 65 hours were the next generation of newly hatched L1 larvae laid by the animals that developed during the time course.

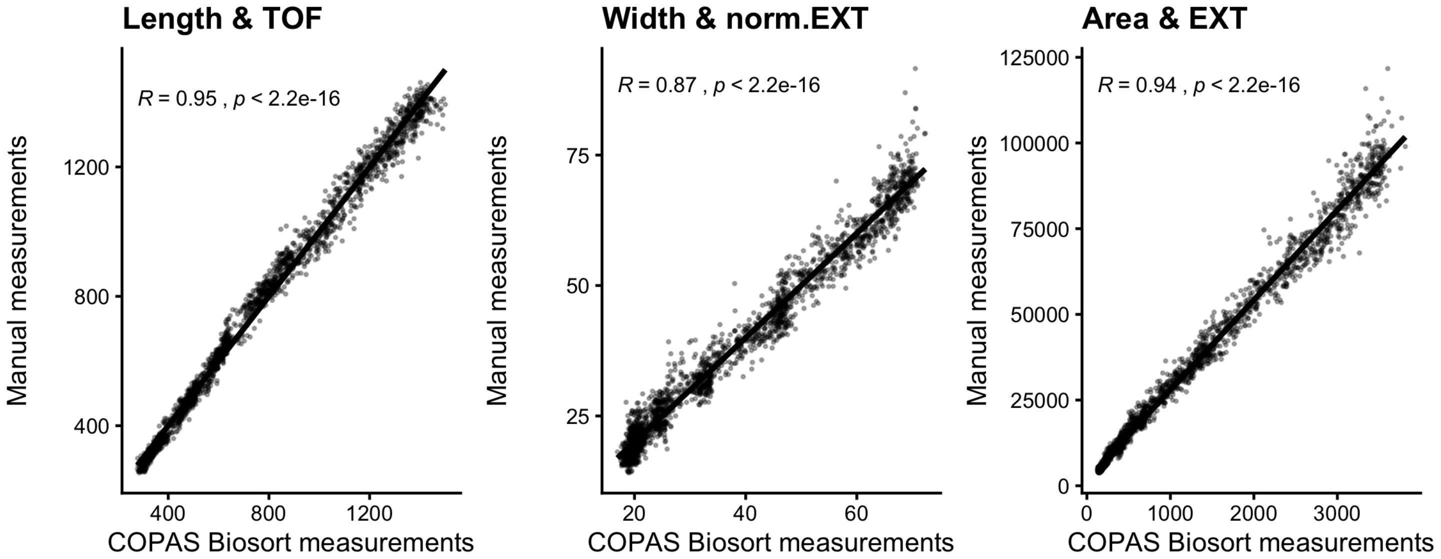


**Figure S2. Correlation analysis of body size measurements.** Manual measurements of animal length, width, and estimated area were compared to COPAS BIOSORT measurements of TOF, norm.EXT, and EXT. Kendall correlation value is shown in each plot.

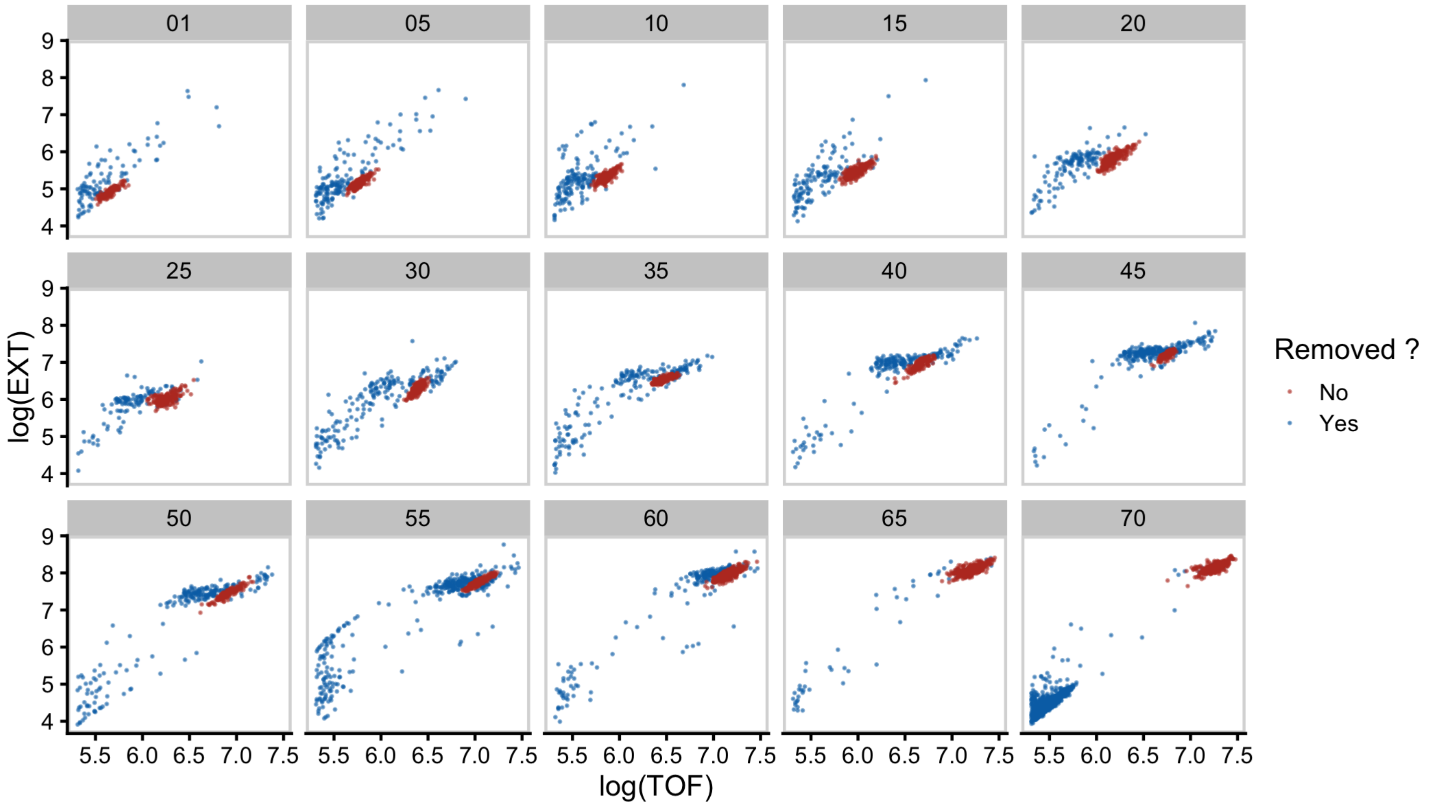


**Figure S3. Mixture modeling of COPAS BIOSORT data was used to prune data.** Mixture models of Gaussian distributions were fit to log transformed animal length (x-axis) and log transformed optical extinction (y-axis). Data from each hour of the experiment was analyzed and processed to remove clusters that did not include animal objects. All replicates were pruned independently; a subset of data from replicate 2 is shown here. Panels indicate experimental hours from which data were taken.

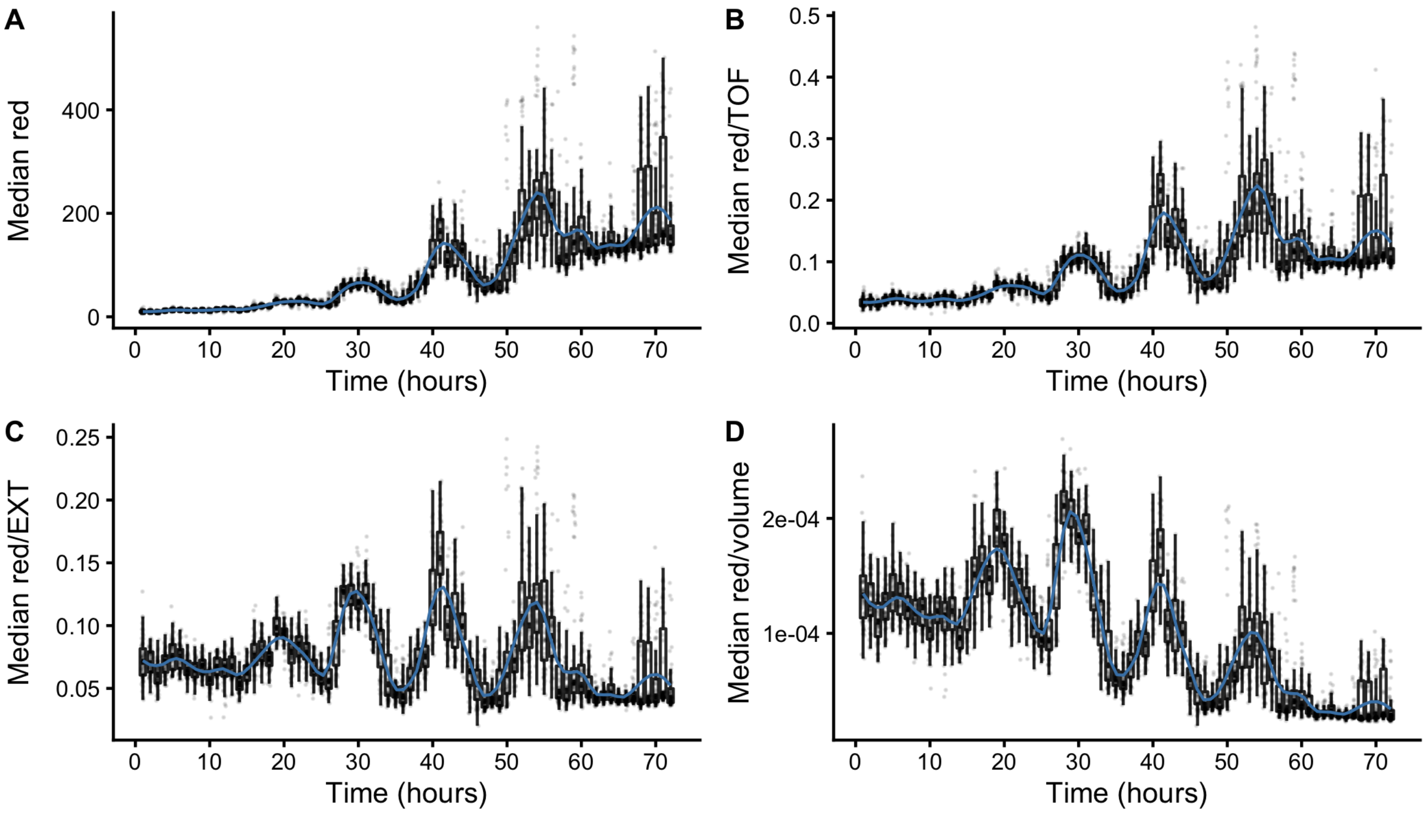


**Figure S4. Fluorescence measurements normalized by body size.** Red fluorescence beads were fed to animals during experimentation and fluorescence data was collected by the COPAS BIOSORT. Fluctuations in fluorescence indicate fluctuations in feeding behavior. Fluorescence data was normalized by body size measurements to account for increases in body size. Dividing fluorescence by area was most successful in normalizing fluorescence dynamics to account for changes in animal size over time.

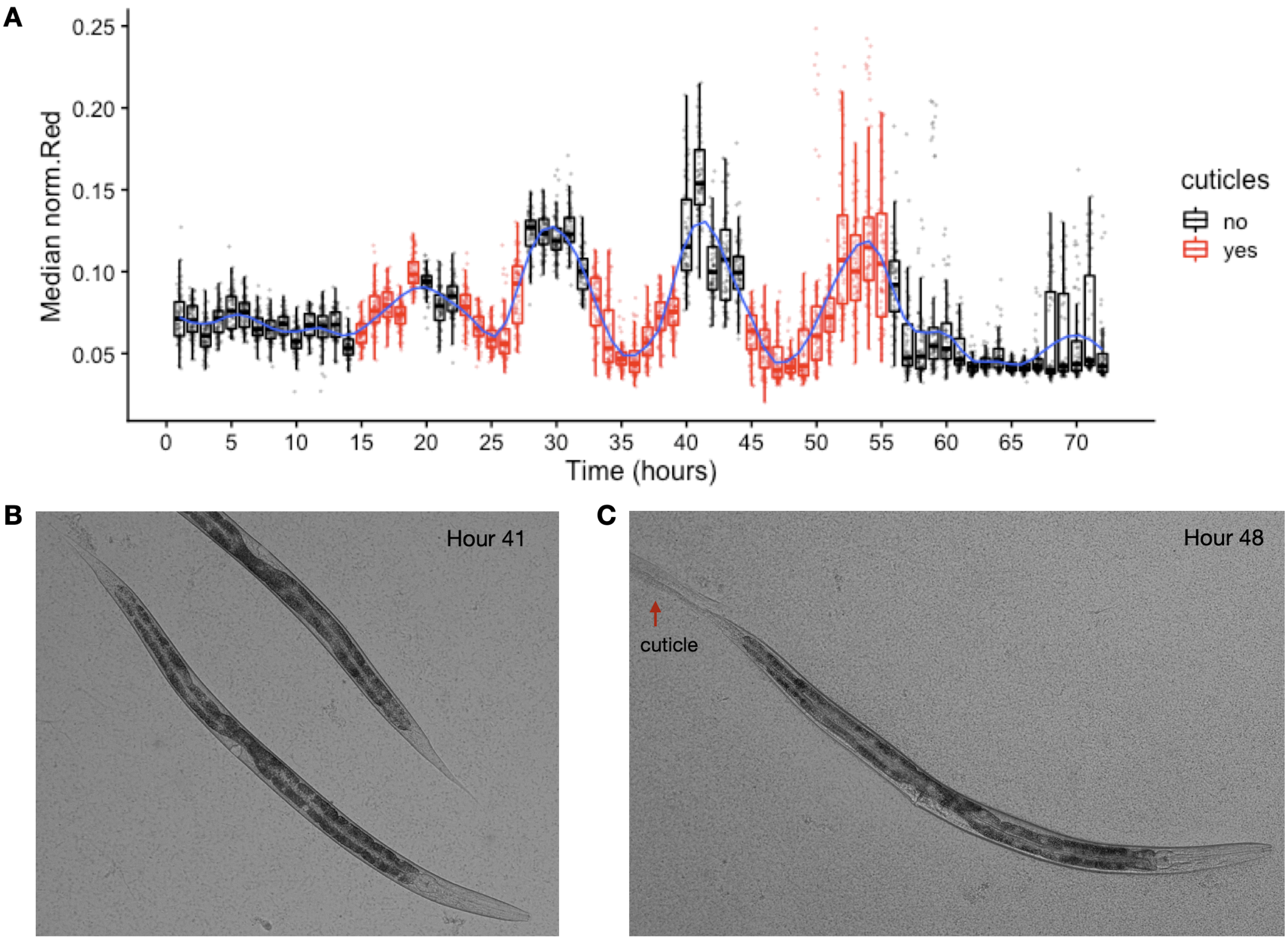


**Figure S5. Cuticles identified during periods of decreased feeding.** Images of wells collected during the experiment were examined for evidence of shed cuticles. (A) Experimental hours where cuticles were identified from images overlap with hours where population feeding behavior is low. Cuticles shed from the L4-Adult molt persisted longer than previous larval stage cuticle debri. (B) Example image of animal without visible cuticle during a period of elevated feeding. (C) Example image of an animal with visible cuticle indicating completion of molt during a period of decreased feeding.

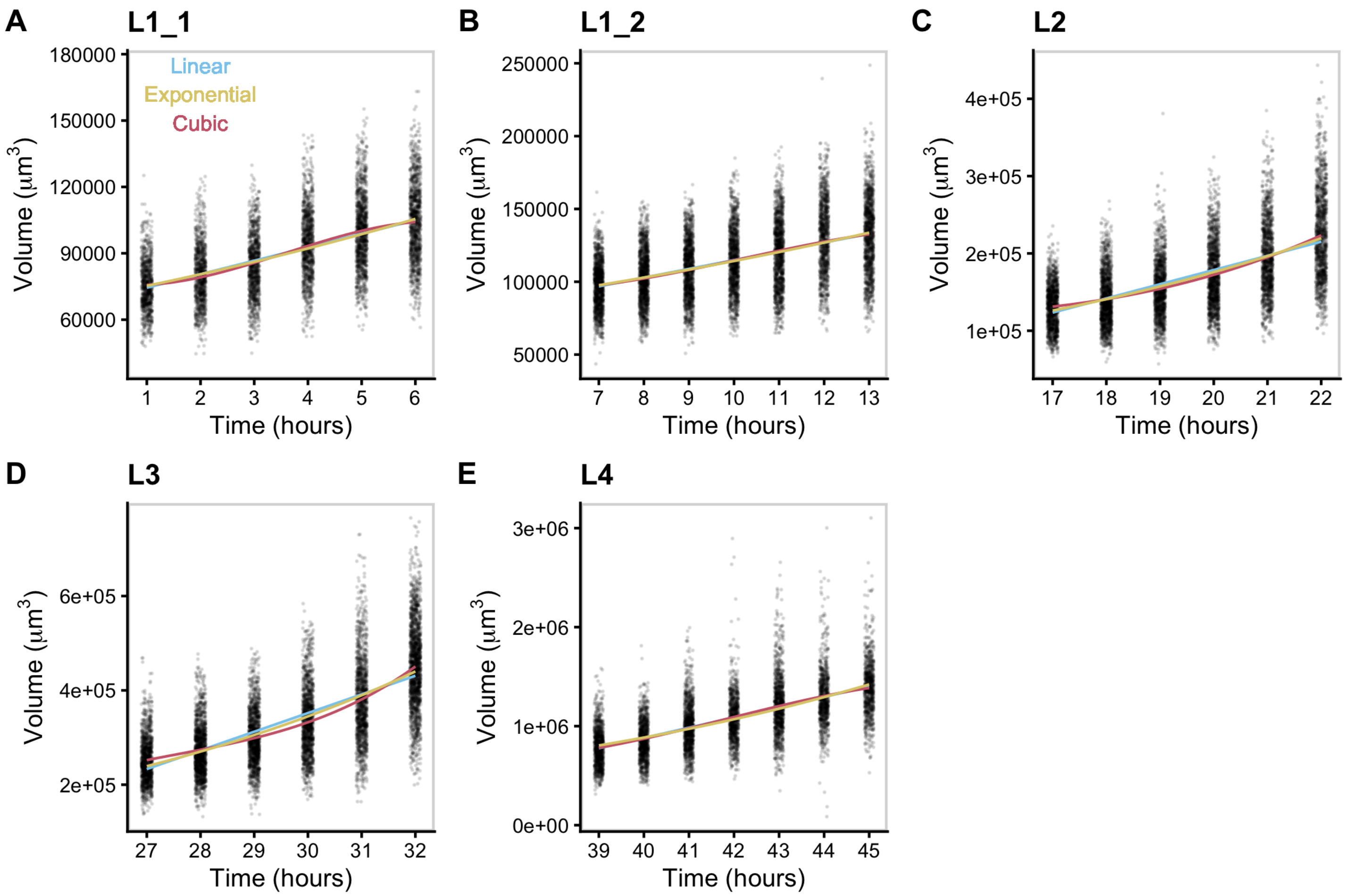


**Figure S6. Volume growth data fit with linear, exponential, and cubic models.** Volume data of individuals in time points defined as growth periods are analyzed for each stage. L1 stage was further separated into two periods to account for the volume dip that occurs mid-stage.

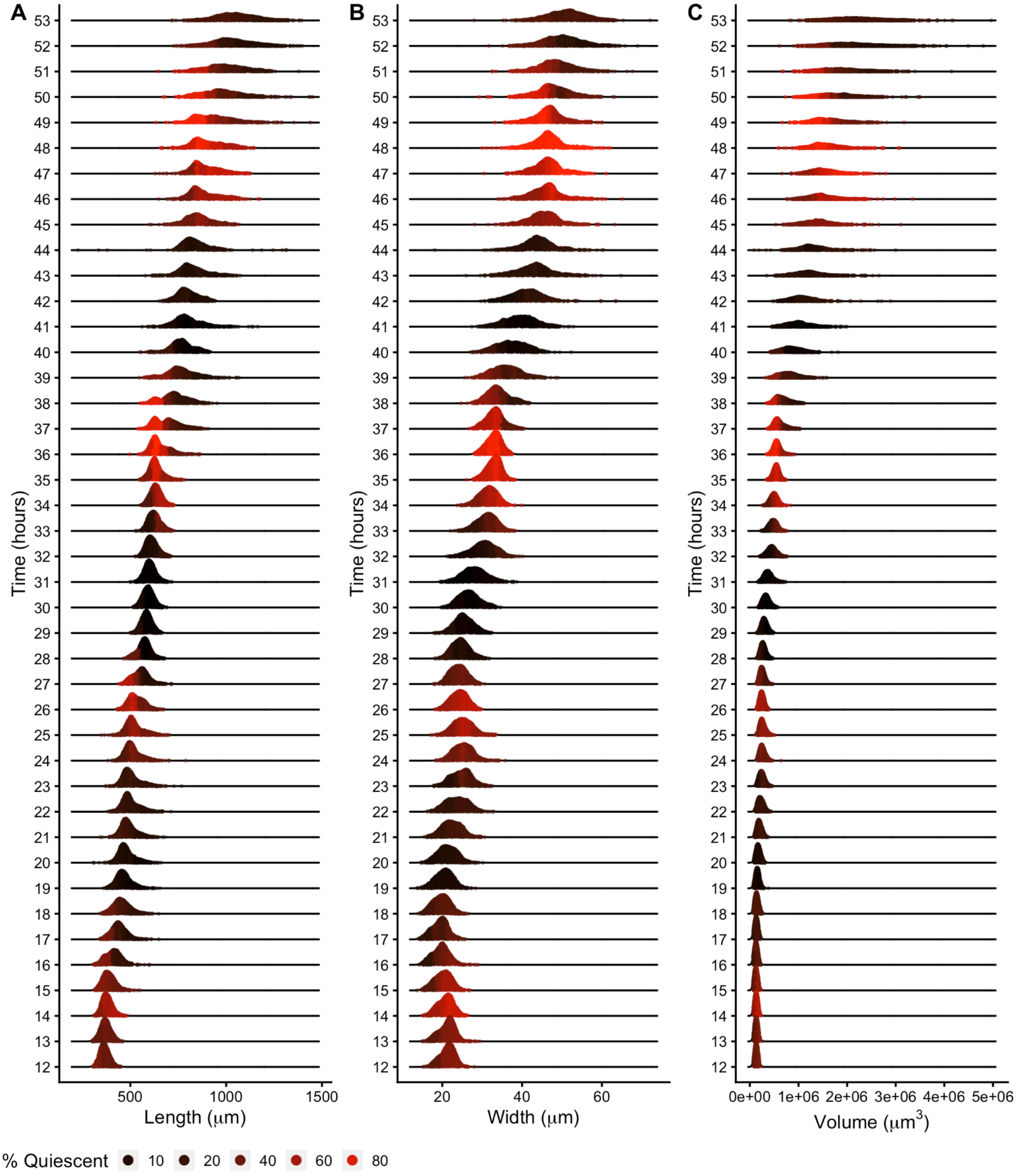


**Figure S7. Density plots of population size dynamics across all larval transitions.** Density curves of length (A), width (B) and volume (C). Curves are divided into five quantiles and colored by the percentage of quiescent animals present within that quantile. Molts are estimated to occur at experimental hours 14, 25, 36, and 48 (see Methods).

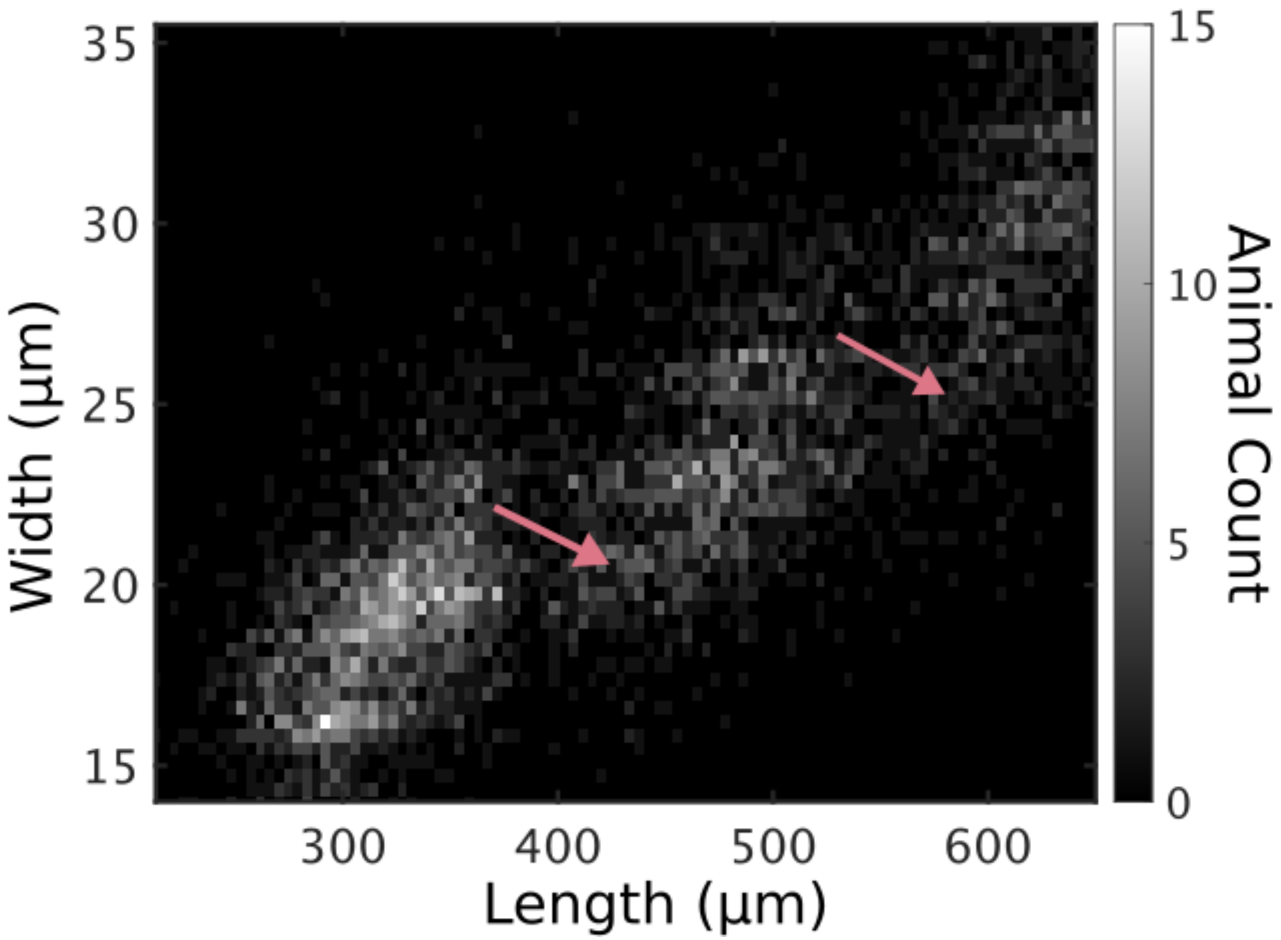


**Figure S8. Animals in all replicates, measured from images.** Animal length and width over *C. elegans* development captured from image data. Higher noise levels in these measurements preclude accurate regressions to individual larval stages. Length jumps and width dips are still apparent. Compare with Fig 6**.**

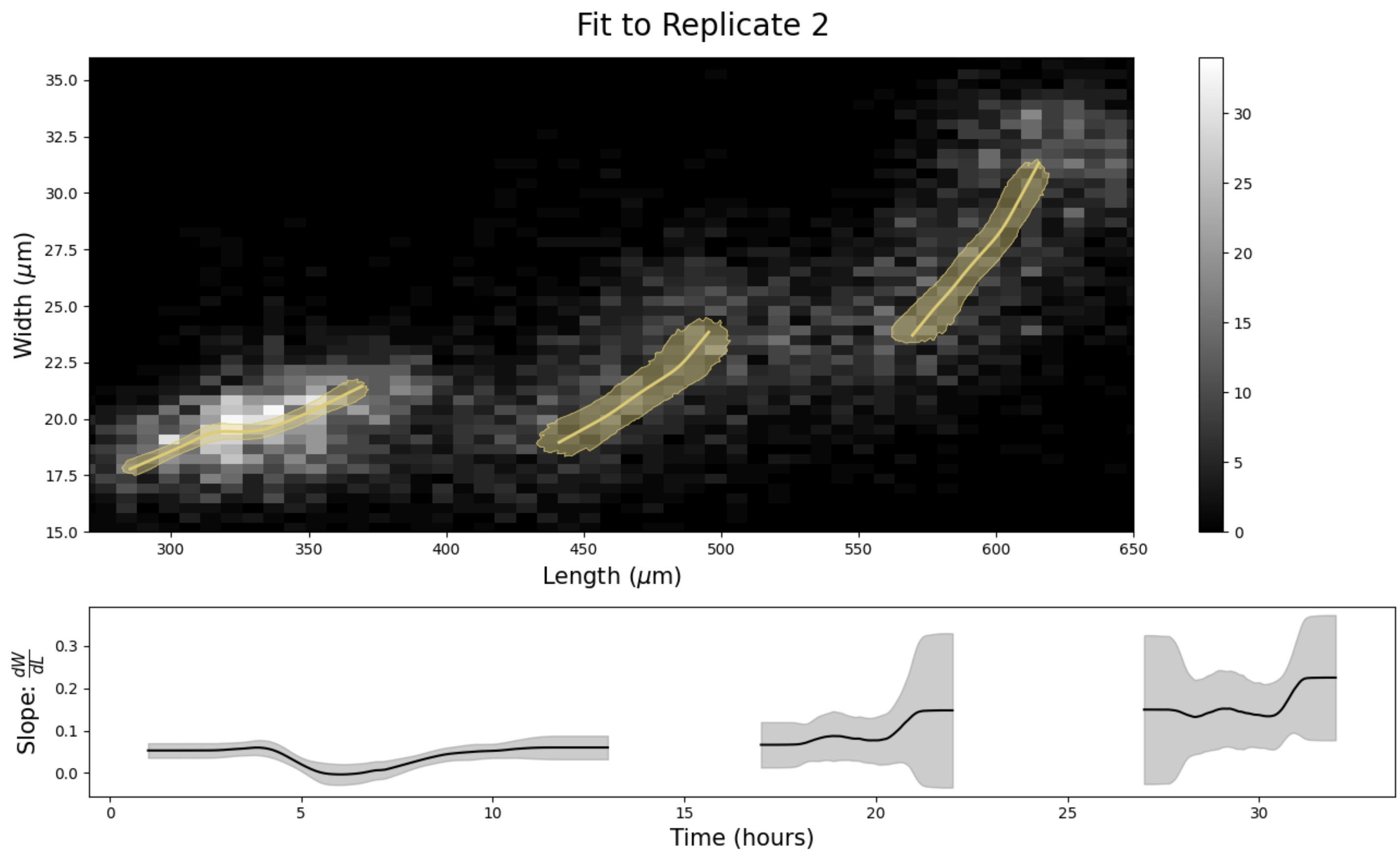


**Figure S9. Stretcher model analysis of replicate 2 COPAS BIOSORT data for different stage thresholding.** Compare to Fig 5**.** Larval hours were defined by taking the ceiling of the lower boundary and the floor of the upper boundary. This rounding method for larval stage definition demonstrates the sensitivity of the analysis to edge effects. The unexpected step in the L2 larval stage (Fig 5) was significantly reduced with this rounding method.

**Table S1. Results of analysis of variance models fit to COPAS BIOSORT data.** Analysis of variance tests were used to quantify the amount of variance in our data contributed by the sampling technique. The sampling technique involved unbiased sampling of animals from six replicate populations and subsequent distribution into multiple wells of a microtiter plate for analysis. We quantified the amount of variance contributed by replicate and well. We find that the variance explained by well is nearly negligible whereas replicate contributes minor variance in some measurements. Given this information, we deem the generated summary statistics an appropriate representation of the population.

**Response = Norm.Red**

| *Terms* | *Df* | *Sum Sq* | *Mean Sq* | *F value* | *Pr(>F)* | *% Var Explained* |
|---|---|---|---|---|---|---|
| hour | 1 | 439.46 | 439.46 | 217762.01 | 0 | 54.34 |
| replicate | 6 | 165.82 | 27.64 | 13694.48 | 0 | 20.51 |
| well | 10 | 0.32 | 0.03 | 15.62 | 0 | 0.04 |
| Residuals | 100619 | 203.06 | 0 | NA | NA | 25.11 |

**Response = Length**

| *Terms* | *Df* | *Sum Sq* | *Mean Sq* | *F value* | *Pr(>F)* | *% Var Explained* |
|---|---|---|---|---|---|---|
| hour | 1 | 86190107879 | 86190107879 | 8223506 | 0 | 98.4 |
| replicate | 6 | 349438944 | 58239824 | 5557 | 0 | 0.4 |
| well | 10 | 834970 | 83497 | 8 | 0 | 0 |
| Residuals | 100619 | 1054582098 | 10481 | NA | NA | 1.2 |

**Response = Width**

| *Terms* | *Df* | *Sum Sq* | *Mean Sq* | *F value* | *Pr(>F)* | *% Var Explained* |
|---|---|---|---|---|---|---|
| hour | 1 | 209090438.47 | 209090438.47 | 6950760.3 | 0 | 97.99 |
| replicate | 6 | 1266985.87 | 211164.31 | 7019.7 | 0 | 0.59 |
| well | 10 | 1495.32 | 149.53 | 4.97 | 0 | 0 |
| Residuals | 100619 | 3026786.99 | 30.08 | NA | NA | 1.42 |

**Response = Volume**

| *Terms* | *Df* | *Sum Sq* | *Mean Sq* | *F value* | *Pr(>F)* | *% Var Explained* |
|---|---|---|---|---|---|---|
| hour | 1 | 621712474191714688 | 621712474191714688 | 843342 | 0 | 84.08 |
| replicate | 6 | 43356968850687072 | 7226161475114512 | 9802 | 0 | 5.86 |
| well | 10 | 152458715405862 | 15245871540586 | 21 | 0 | 0.02 |
| Residuals | 100619 | 74176459811167296 | 737201321929 | NA | NA | 10.03 |

**Table S2. Model fit criteria used to assess candidate growth models.** To determine the level of support for each model, the candidate model with the smallest raw AIC/BIC was identified and compared to other AIC/BIC values. If the delta value was greater than 6, the model with the smallest AIC/BIC value was denoted as the best model. If the delta value was less than 6 but greater than 2, the model with the smallest AIC/BIC value was determined to likely be the best model. If the delta value was less than 2, we are unable to distinguish the model of best fit.

| | $\Delta AIC$ | | | $\Delta BIC$ | | | | |
|---|---|---|---|---|---|---|---|---|
| Stage | Linear | Exponential | Cubic | Linear | Exponential | Cubic | Best model by AIC | Best model by BIC |
| L1_1 | 17 | 21 | 0 | 4 | 9 | 0 | Cubic | Likely Cubic |
| L1_2 | 2 | 4 | 0 | 0 | 2 | 12 | Can't distinguish | Can't distinguish |
| L2 | 142 | 43 | 0 | 128 | 28 | 0 | Cubic | Cubic |
| L3 | 374 | 145 | 0 | 360 | 131 | 0 | Cubic | Cubic |
| L4 | 4 | 44 | 0 | 0 | 40 | 10 | Likely Cubic | Linear |

### S5 File. Model Derivations

#### Bootstrapping Algorithm

To calculate a robust regression of the measured COPAS BIOSORT data we bootstrap the regression using case-resampling (Davison and Hinkley 1997, pages 261-266). Each iteration of the algorithm (**Algorithm S1**) involves resampling the data of interest with replacement and maintaining the size of the sample. At each iteration the lokern regression is calculated for Red, Length, and Width data. Additionally any desired function (for example, volume, pumping frequency times pharynx fraction, or derivatives) of these regressions is calculated at each iteration. Regressions at each iteration are saved and the mean and variance of these regressions at each regression time point are used to determine the statistics of the regression.

**Algorithm S1** Regression Bootstrapping with Case Resampling

```
Ndata = # animals in sample;
iterations = # resamplings;

for i = 0 to iterations do
   Resample Ndata points with replacement. Collect Red, Length, Width data;

   Apply lokern regression to resampled Red, Length, Width;

   Calculate desired functions of Red, Length, Width regressions;

   Save Red, Length, Width, and combined regressions;
end for

Calculate Standard deviation at each regression time point of saved regressions;
```

#### Derivation of eating model

We begin by defining the instantaneous rate of food intake as a function of the flow rate of media through the buccal cavity and the cross sectional area of the buccal cavity.

$$\frac{dV_{food}}{dt} = A_{buccal} \tag{S1}$$

We then make the assumption that the uptake of media fills the pharyngeal lumen we have

$$\frac{dV_{food}}{dt} = C\frac{dV_{lumen}}{dt} \tag{S2}$$

We average both sides of the equation under the assumption that the pumping period is significantly shorter than the time scale of growth

$$\frac{1}{T}\int_0^T \frac{dV_{food}}{dt}dt = \frac{1}{T}\int_0^T CA_{buccal} \tag{S3}$$

The integral on the right hand side is the total food intake during a single pumping period.

$$\frac{1}{T}\int_0^T \frac{dV_{food}}{dt}dt = \frac{1}{T}\Delta V_{food} \tag{S4}$$

We then take into account that food is not transported to the gut in the same step as its uptake. Thus the total food intake during a single pump can be calculated by the amount of food that fills the fully opened pharyngeal lumen

$$\frac{1}{T}\int_0^T \frac{dV_{food}}{dt}dt = \frac{C}{T}V_{lumen:max} \tag{S5}$$

We then replace the average on the right hand side with the average food intake rate over the pumping period. For simplicity and because we will deal entirely with the average food intake rate, we do not use a different notation for this average rate.

$$\frac{dV_{food}}{dt} = \frac{C}{T}V_{lumen:max} = Cf(t)V_{lumen:max} \tag{S6}$$

#### Transformation of sorter measurements to volume units

To utilize sorter measurements and convert them to meaningful units we define a linear transformation from the correlation plots (S1 Fig)

$$L = a_1 TOF + b_1 \tag{S7}$$

$$W = a_2 norm.EXT + b_2. \tag{S8}$$

Using Equations (S7) and (S8) we can approximate the volume of any object that passes through the sorter by the expression

$$V = \frac{\pi}{4}(a_1 TOF + b_1)(a_2 norm.EXT + b_2)^2. \tag{S9}$$

#### Defecation analysis

We use the defecation results found in (Liu and Thomas 1994) to determine if red fluorescent measurements can be used as a proxy for food intake rate as opposed to the instantaneous food volume in the gut. Defecation in adults happens very regularly, with a period of $T_d = 45 \pm 3s$, with $h_d = 43 \pm 10\%$ of their intestinal volume being expelled each time. The volume expelled is well mixed. Defining $V_f$ as the current amount of food in the nematode gut. We can use conservation of mass to state that the rate of change in the amount of food in the gut is equal to the rate of food intake through eating less the defecation rate and the rate at which food volume is metabolized into cell products:

$$\frac{dV_f}{dt} = \left.\frac{dV_f}{dt}\right|_{eating} - \left.\frac{dV_f}{dt}\right|_{defecating} - \left.\frac{dV_f}{dt}\right|_{metabolized} \tag{S10}$$

Using red fluorescence as a proxy for food intake we can ignore the metabolism term as the fluorescent beads are not metabolized.

$$\frac{dV_f}{dt} = \left.\frac{dV_f}{dt}\right|_{eating} - \left.\frac{dV_f}{dt}\right|_{defecating} \tag{S11}$$

Equation (S11) states that the rate of change of the volume of fluorescent beads in the gut is equal to the difference between the intake of red fluorescent beads minus the defecation rate of red fluorescent beads. Using the results of (Liu and Thomas 1994) for the second term, and defining $V_{f:max}$ as the volume of food in the gut just prior to defecation, $h_d$ as the fraction of food expelled during a single defecation cycle, and $T_d$ as the period of defecation. We average Equation (S11) over short time periods to remove the pumping and defecation period oscillations.

$$\frac{dV_{red:max}}{dt} = \left.\frac{dV_red}{dt}\right|_{eating} - \frac{V_{red:max}h_d}{T_d} \tag{S12}$$

On the left hand side, the instantaneous rate of the gut red fluorescence in Equation (S11) is replaced by the rate of change of the maximum or "full" gut fluorescence. The second term on the right hand side of Equation (S11) has been replaced by the average defecation rate over a cycle calculated by multiplying the full gut fluorescence by the fraction expelled and dividing by the defecation period. We solve Equation (S12) for the average eating rate.

$$\left.\frac{dV_{red}}{dt}\right|_{eating} = \frac{dV_{red:max}}{dt} + \frac{V_{red:max}h_d}{T_d} \tag{S13}$$

We take the local regression of the red fluorescence to determine $V_{red:max}(t)$. This value is plugged into the second term on the right hand side of Equation (S13) and its derivative is used to approximate the first term on the right hand side of Equation (S13). We take the adult values of $h_d$ and $T_d$ (Liu and Thomas 1994) as a first approximation. Figure (S6) demonstrates a comparison of the red fluorescence and the red intake rate with defecation taken into account at the constant adult rates and quantities. We have scaled both the pink curve denoting red fluorescence and the blue curve denoting red intake rate by their maximum. This scaling allows us to see that the two curves are only a multiplicative factor apart up to error bars. This allows us to use the red measurement as a proxy for both red and food intake rates.

#### Food Allocation Breakdown Calculation

We calculate an example of resource allocation breakdown. We begin with the expression in Equation (8) of the main text relating food intake rate and animal growth rate and repeated here:

$$\frac{dV_{worm}}{dt} = \eta(t)\alpha(t)\frac{dV_{food}}{dt} \tag{S14}$$

Here $\eta(t)$ is metabolic efficiency and is characteristic of the nematodes and food source used. $\alpha(t)$ is the fraction of total ingested food allocated towards growth.

We make an additional assumption that food needed to maintain life is proportional to animal volume:

$$\frac{dV_{food:maint}}{dt} = \beta V_{worm}(t) \tag{S15}$$

Here we define $\frac{dV_{food:maint}}{dt}$ as the rate of food intake required to maintain life and $\beta$ is the constant of proportionality that relates animal size to required maintenance food levels. We assume that $\beta$ is a constant over developmental time and depends on the food source available.

From the analysis from the previous section we can use red fluorescence as a proxy for food intake rate such that

$$\frac{dV_{food}}{dt} \propto Red \tag{S16}$$

$$\frac{dV_{worm}}{dt} \propto \eta(t)\alpha(t)Red \tag{S17}$$

$$\eta(t)\alpha(t) \propto \left(\frac{dV_{worm}}{dt}\right)/Red \tag{S18}$$

The constant of proportionality in Equation (S18) is unknown due to the unknown relationship between food volume and red fluorescence. We make the assumption that metabolic efficiency is constant over time and any variation in the value calculated by Equation (S18) is due to changes in $\alpha(t)$, the allocation of food toward growth.

To calculate an estimate for the breakdown of food allocation, we assume a set of constants of proportionality for Equations (S15) and (S18). We assume that at the time at which the $\eta(t)\alpha(t)$ curve is at its maximum ( hour 40), 10% of food resources are allocated towards maintenance, 70% (low growth estimate) or 90% (high growth estimate) are allocated towards growth, and the remainder is allocated to other metabolic processes. The resulting scaling factors are applied over developmental time to calculate an estimated food allocation breakdown and presented in Figure (8) of the main text.

### Derivation of Stretcher Model

We model the cuticle as a thin walled pressure vessel made of orthotropic, linear materials to capture the relationship between how much the cuticle stretches and the force applied to the cuticle. The relationship between the amount of stretch and the amount of applied pressure is described by the matrix:

$$\begin{bmatrix} \varepsilon_L \\ \varepsilon_{Circ} \end{bmatrix} = \begin{bmatrix} \frac{1}{E_L} & \frac{-v_{cl}}{E_c} \\ \frac{-v_{lc}}{E_L} & \frac{1}{E_c} \end{bmatrix} \begin{bmatrix} \sigma_L \\ \sigma_{Circ} \end{bmatrix} \tag{S19}$$

Here $E_L$ and $E_C$ are the Young's modulus in the length and circumferential direction, and $v_{cl}$ and $v_{lc}$ are the appropriate Poisson's ratios. These material properties can be measured experimentally. Normalized stretch, $\varepsilon$, is defined as the change in size normalized by the initial size of a cuticle in length (L) and circumference (Circ) (**Equations S20 - S21**) . Here $\sigma$ is the normalized force applied along the length and circumferential directions of the cuticle (**Equations S23 - S24**).

$$\varepsilon_L = \frac{\Delta L}{L_0} \tag{S20}$$

$$\varepsilon_{Circ} = \frac{\Delta Circ}{Circ_0} \tag{S21}$$

Here $L_0$ and $Circ_0$ are the length and circumference of the cuticle at the onset of stretch, or in other words at the start of a larval stage. Experimentally we measure width, not circumference, but the circumference of a circle is proportional to the width of the circle, so we can replace the circumference with the measured width:

$$\frac{\Delta Circ}{Circ_0} = \frac{\pi \Delta W}{\pi W_0} = \frac{\Delta W}{W_0} \tag{S22}$$

By approximating *C. elegans* as cylindrical, and assuming the only force working on the cuticle is isotropic internal pressure, we can determine the normalized force in terms of pressure and geometric properties:

$$\sigma_L = \frac{r}{2t}\Delta p \tag{S23}$$

$$\sigma_{Circ} = \frac{r}{t}\Delta p \tag{S24}$$

Where r is the radius of the cylinder and t is the thickness of the cuticle. We can now rewrite Equation (S19) in terms of measurable quantities (length and width) by multiplying out the matrices making the appropriate substitutions.

$$\Delta L = \frac{L_0 r}{t}\left(\frac{1}{2E_L} - \frac{v_{cl}}{E_c}\right)\Delta p = a_L \Delta p \tag{S25}$$

$$\Delta W = \frac{W_0 r}{t}\left(\frac{1}{E_C} - \frac{v_{lc}}{E_L}\right)\Delta p = a_W \Delta p \tag{S26}$$

Here we compress the coefficients fixed by material and geometric properties into one constant

$$a_L = \frac{L_0 r}{t}\left(\frac{1}{2E_L} - \frac{v_{cl}}{E_c}\right) \tag{S27}$$

$$a_W = \frac{W_0 r}{t}\left(\frac{1}{E_c} - \frac{v_{lc}}{E_L}\right) \tag{S28}$$

### Stretcher Slope Calculations

The Stretcher model predicts a constant ratio between stretch in width, $\Delta W$, and stretch in length, $\Delta L$. These measures of stretch are changes in the length and width measurements over some period of time. To see how the ratio of $\frac{\Delta W}{\Delta L}$ changes throughout a larval stage, we need to measure the stretch instantaneously. To do this we take advantage of the derivative approximation

$$\Delta W \approx \frac{dW}{dt}\Delta t \tag{S29}$$

$$\Delta L \approx \frac{dL}{dt}\Delta t \tag{S30}$$

Equations (S29, S30) are combined in the ratio found in Equation (3) to give:

$$\frac{\Delta W}{\Delta L} \approx \frac{W'(t)}{L'(t)} \tag{S31}$$

Here $W' = \frac{dW}{dt}$ and $L' = \frac{dL}{dt}$. We differentiate the time series from the local regressions of length and width data. The ratio of derivatives gives the instantaneous stretch ratio as it changes over time. Due to the numerical difficulty of calculating both derivatives and ratios with accuracy, we expect large error bars for this ratio. To estimate the size of error bars we apply a first order error propagation formula to Equation (S31).

$$\sigma^2_{Ratio} \approx \left|\frac{\partial Ratio}{\partial (W')}\right|^2 \sigma^2_{(W')} + \left|\frac{\partial Ratio}{\partial (L')}\right|^2 \sigma^2_{(L')} + 2\frac{\partial Ratio}{\partial (W')}\frac{\partial Ratio}{\partial (L')}\sigma_{(W')(L')} \tag{S32}$$

$$\sigma^2_{Ratio} \approx \left|\frac{1}{L'}\right|^2 \sigma^2_{(W')} + \left|\frac{-(W')^2}{(L')^3}\right|^2 \sigma^2_{(L')} - 2\frac{W'}{(L')^3}\sigma_{(W')(L')} \tag{S33}$$

The variance in Equation (S33) is calculated for each time point in the regressions of length and width. We resample the data for each larval stage 2,000 times with replacement (see Methods for larval stage determination). For each of these resampled sets of data, we numerically differentiate the length and width regression to determine an estimate of the length and width derivatives over time. At each time point the mean regression length and width derivatives are used in place of $W'$ and $L'$. The covariance matrix for the length and width derivatives is calculated at each time point using the 2,000 resampled regressions in the bootstrap analysis and used as the variance terms in Equation (S33).
